# FANCM regulates repair pathway choice at stalled replication forks

**DOI:** 10.1101/2020.10.29.357996

**Authors:** Arvind Panday, Nicholas A. Willis, Rajula Elango, Francesca Menghi, Erin E. Duffey, Edison T. Liu, Ralph Scully

## Abstract

Conservative repair of stalled replication forks is important for the maintenance of a stable genome. However, the mechanisms that regulate repair pathway “choice” at stalled mammalian forks remain poorly understood. The Fanconi anemia complementation group M gene, *FANCM*, encodes a multi-domain scaffolding and motor protein that interacts with several distinct repair protein complexes at stalled forks. Here we use a chromosomally integrated reporter of stalled fork repair, in combination with defined mutations engineered within the endogenous *Fancm* gene in primary mammalian cells, to study how *Fancm* regulates stalled fork repair. We identify separation-of-function *Fancm* mutants, which reveal that distinct repair functions of FANCM are enacted by modular, molecularly separable scaffolding domains. These findings define FANCM as a key mediator of repair pathway choice at stalled replication forks and reveal its molecular mechanism. Notably, a mutation that inactivates the ATPase function of FANCM disables all FANCM-mediated repair functions and appears to “trap” FANCM at stalled forks. We find that *Fancm* null cells do not survive genetic inactivation of *Brca1*. This synthetic lethal interaction is recapitulated in *Fancm* ATPase-defective mutants. The ATPase function of FANCM may therefore represent a promising “druggable” target for therapy of *BRCA1* mutant cancers.

## Introduction

Genomic instability underlies the pathogenesis of many human diseases, ranging from rare hereditary developmental syndromes to common diseases such as cancer (Ciccia and Elledge, 2010; Taylor et al., 2019). A major threat to genome stability arises when replication forks stall at sites of DNA damage or abnormal DNA structure (Cortez, 2019; Zeman and Cimprich, 2014). Eukaryotic replication fork stalling activates S-phase checkpoint signaling, triggers fork remodeling and exposes the stalled fork to a range of possible repair activities (Neelsen and Lopes, 2015; Quinet et al., 2017; Rickman and Smogorzewska, 2019). In vertebrates, conservative repair of stalled forks requires the homologous recombination (HR) pathway, which mediates protective fork-remodeling in addition to canonical repair (Duxin and Walter, 2015; Schlacher et al., 2011; Scully et al., 2019). Alternative, error-prone repair pathways include aberrant DNA end joining and replication restart mechanisms (Adamo et al., 2010; Pace et al., 2010; Willis et al., 2017). The “choice” between conservative and error-prone repair of stalled forks has a major bearing on the stability of the genome, but the mechanisms underlying repair pathway choice at the stalled fork remain poorly understood.

Stalled fork repair is defective in humans with Fanconi anemia (FA)—a rare, recessive genetic disorder associated with childhood anemia and increased cancer incidence (Niraj et al., 2019). Cells derived from FA patients reveal spontaneous genomic instability and are abnormally sensitive to DNA interstrand crosslinking agents (ICLs), such as mitomycin C, as well as to DNA-protein crosslinking agents such as aldehydes (Langevin et al., 2011; Rosado et al., 2011). Twenty-two distinct FA genes have been identified to date, defining a pathway that includes the hereditary breast and ovarian cancer (HBOC) predisposition genes, *BRCA1/FANCS* and *BRCA2/FANCD1*, as well as other general HR genes such as *RAD51* (Niraj et al., 2019; Prakash et al., 2015). A small proportion of HBOC risk in the human population is accounted for by monoallelic germline mutation of FA genes other than *BRCA1* and *BRCA2*. These genes include *Rad51C/FANCO* and *FANCM* (Bogliolo et al., 2018; Castéra et al., 2018; Catucci et al., 2018; Figlioli et al., 2020; Neidhardt et al., 2017; Peterlongo et al., 2015).

ICL processing by the FA pathway is activated by bidirectional replication fork arrest at the ICL, followed by replisome disassembly and asymmetrical fork reversal, with recruitment of the FANCD2/FANCI heterodimer (Amunugama et al., 2018; Deans and West, 2011; Knipscheer et al., 2009; Long et al., 2011; Raschle et al., 2008). FANCD2/FANCI is monoubiquitinated by the FA “core” complex, comprised of FANC A, B, C, E, F, G and L—FANCL being the subunit with E3 ubiquitin ligase activity (Kim and D’Andrea, 2012; Meetei et al., 2003). Ubiquitinated FANCD2, in turn, recruits the SLX4 nuclease scaffold, which mediates dual incisions of one sister chromatid to generate an “unhooked” ICL on one sister and a two-ended DSB on the other (Hodskinson et al., 2014; Klein Douwel et al., 2014; Zhang and Walter, 2014).

Mammalian *FANCM* (Fanconi anemia complementation group M) encodes a large scaffolding and motor protein that is part of a family of conserved DEAH-type DNA-dependent ATPases and dsDNA translocases related to the Archaeal Hef protein (Meetei et al., 2005; Nandi and Whitby, 2012; Prakash et al., 2005; Sun et al., 2008; Whitby, 2010; Xue et al., 2015). Although Hef is both a helicase and a nuclease, the FANCM nuclease domain is inactive in higher eukaryotes. FANCM homologs display branch migration, D loop dissociation, replication fork reversal and 3’-5’ DNA helicase activity, the last of these being absent from the mammalian protein (Gari et al., 2008a; Gari et al., 2008b; Meetei et al., 2005; Sun et al., 2008; Whitby, 2010; Xue et al., 2015; Zheng et al., 2011). At the cellular level, FANCM homologs contribute to the non-crossover “synthesis-dependent strand annealing” (SDSA) pathway of HR (Paques and Haber, 1999; Prakash et al., 2009; Sun et al., 2008). FANCM homologs suppress meiotic and mitotic crossing over (Crismani et al., 2012; Knoll et al., 2012; Romero et al., 2016) including sister chromatid exchange (SCE), a cytological measure of crossing over (Bakker et al., 2009; Deans and West, 2009; Rosado et al., 2009). FANCM ATP hydrolysis mutants are defective for ICL resistance, indicating that FANCM motor function contributes to stalled fork repair (Nandi and Whitby, 2012; Xue et al., 2008).

FANCM binds FAAP24 and two <u>M</u>ajor <u>H</u>istone <u>F</u>old proteins MHF1 and MHF2 (Ciccia et al., 2007; Singh et al., 2010; Tao et al., 2012; Yan et al., 2010). This complex facilitates FANCM recruitment to stalled forks and ATR-dependent S phase checkpoint responses (Collis et al., 2008). The FANCM MM1 domain binds the FA core complex *via* direct interaction with FANCF, while the FANCM MM2 domain binds the Bloom’s syndrome helicase (BLM)-TOP3A-RMI1-RMI2 (BTR) complex *via* direct interaction with RMI1 (Deans and West, 2009) (**Figure 1A**). Both FANCM ΔMM1 and MM2 mutants were found to be defective for ICL-resistance and for SCE suppression (Deans and West, 2009). It is currently unclear how individual biochemical functions of FANCM are connected to specific pathways of stalled fork repair. Notably, all known *FANCM* mutations in HBOC are predicted to truncate and destabilize the gene product, providing limited information regarding the relationship between individual FANCM domains and HBOC risk (Figlioli et al., 2020).

**Figure 1.**
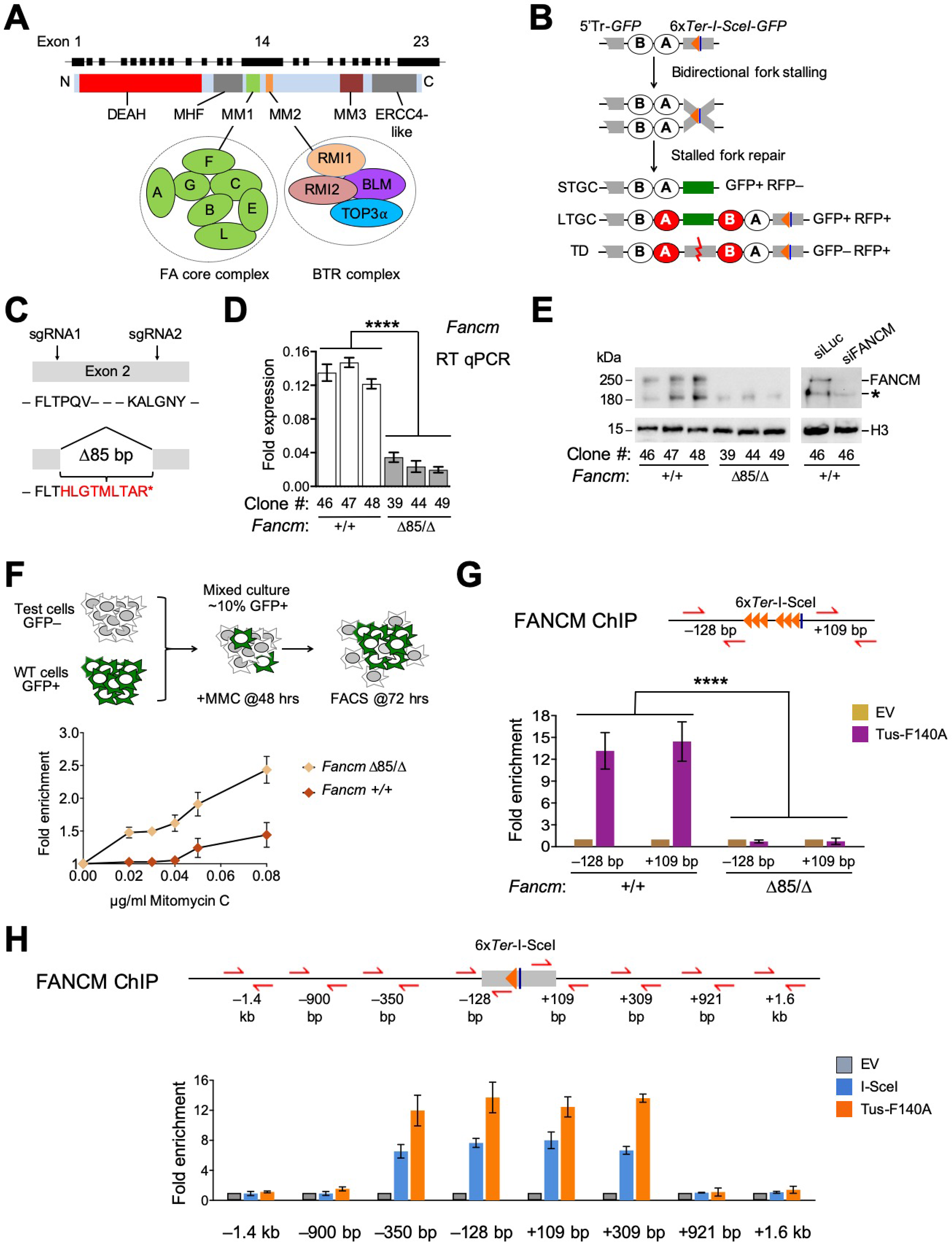
FANCM is recruited to Tus/*Ter*-stalled mammalian replication forks *in vivo*. See also Figure S1. **A.** Schematic of FANCM protein with gene structure above. DEAH: helicase domain. Interactions of MM1 domain with Fanconi Anemia core complex and MM2 domain with the BTR complex are shown. **B.** 6x*Ter*-HR reporter and repair products of Tus/*Ter*-induced fork stalling. Grey boxes: mutant *GFP* alleles. Orange triangle: *6xTer* array. Blue line: *I-SceI* restriction site. Ovals A and B: artificial 5’ and 3’ *RFP* exons. Red ovals: wild-type *RFP* coding exons. STGC/LTGC: short/long tract gene conversion. TD: tandem duplication. Red zig-zag line: non-homologous TD breakpoint. **C.** Plan of 85 bp frame shift *Fancm*^Δ85^ allele. Letters indicate open reading frame (ORF). Red letters: frame-shift ORF with premature stop codon (*). **D.** RT qPCR analysis of *Fancm* mRNA normalized to *Gapdh* mRNA using the 2^−ΔCT^ method in three independent experiments (n=3). Error bars, standard deviation. ****: P < 0.0001 by oneway ANOVA. **E.** Left panel: FANCM immunoblot of chromatin fraction in Fancm^+/+^ and *Fancm*^Δ85/Δ^ clones. H3: Histone H3 loading control. Right panel: loss of FANCM band in siFANCM-treated samples. siLuc: control siRNA to Luciferase. *: background band. **F.** Proliferative competition assay in presence of Mitomycin C (MMC), measuring enrichment of GFP^+^ wild type *vs*. GFP^−^ test culture. Data, normalized to 0 μg/mL MMC, shows mean (n=3) and standard deviation. **G.** ChIP analysis of FANCM at Tus/*Ter* RFB. Upper panel: Schematic of *1xGFP* 6x*Ter*-I-SceI at *Rosa26* (*GFP* sequences not marked). Orange triangles: *Ter* sites. Blue line: I-SceI restriction site. Red half-arrows: qPCR primers. Numbers indicate distance in bp from outer primer to 6x*Ter* array. Lower panel: FANCM ChIP 24 hours after transfection of cells shown with empty vector (EV; gold) or Tus-F140A (purple). Data shows mean of 2^−ΔΔCT^ values (n=3), normalized to EV and *β-Actin* control locus. Error bars: standard deviation. ****: P < 0.0001 by one-way ANOVA. **H.** ChIP analysis of FANCM at *Tus/Ter* RFB or I-SceI-induced DSB (n=3). Schematic of ChIP locus as in panel G. Grey box: 6x*Ter*-I-SceI *GFP* allele. Data obtained and analyzed as in panel G.

The *Saccharomyces cerevisiae FANCM* homolog, *MPH1*, together with the *ScBLM* homolog, *SGS1*, delays the onset of break-induced replication (BIR) in response to an HO-induced DSB (Jain et al., 2016). *ScMPH1* also mediates template-switching during the onset of BIR (Stafa et al., 2014). These observations suggest that FANCM can disassemble BIR intermediates *in vivo*. Mammalian FANCM can also mediate “traverse” of an ICL by the replicative helicase (Huang et al., 2013). FANCM and BLM have both collaborative and distinct roles in the Alternative Lengthening of Telomeres (ALT) pathway, acting concertedly to suppress ALT initiation at stalled forks; BLM additionally supports ALT-related telomere synthesis (Lu et al., 2019; Pan et al., 2017; Silva et al., 2019).

To study stalled fork repair in mammalian cells, we previously adapted the *Escherichia coli* Tus/*Ter* replication fork barrier (RFB) to induce site-specific, bidirectional replication fork stalling and HR at a defined chromosomal locus in mammalian cells (Willis et al., 2014; Willis and Scully, 2016). By placing an array of six 23 bp *Ter* sites within an HR reporter adjacent to a target site for the rare-cutting homing endonuclease I-SceI, we were able to quantify stalled fork repair and, in parallel, conventional DSB repair. We identified three distinct pathways of stalled fork repair (**Figure 1B**): “short tract” gene conversion (STGC), “long tract” gene conversion (LTGC) and the formation of non-homologous tandem duplications (TDs) (Willis et al., 2017).

STGC is a conservative HR outcome, in which limited gene conversion between two mutant heteroalleles of *GFP* (encoding enhanced GFP) converts the cell to GFP^+^. Tus/*Ter*-induced STGC is a product of two-ended HR, suggesting that it is a product of bidirectional fork stalling (Willis et al., 2014; Willis et al., 2017). It is mediated by BRCA1, BRCA2, Rad51 and other canonical HR genes. In contrast to DSB-induced HR, which competes with the second major DSB repair pathway, classical non-homologous end joining (cNHEJ), Tus/*Ter*-induced STGC is unaffected by the status of cNHEJ (Scully et al., 2019; Willis et al., 2018).

LTGC is an error-prone replicative response, potentially related to BIR in yeast (Chandramouly et al., 2013; Saini et al., 2013; Willis et al., 2015). LTGC, which accounts for a minority of HR products in wild type cells, results in an expansion of the repaired sister chromatid and converts the cell to GFP^+^RFP^+^ (**Figure 1B**). Unlike its DSB-induced counterpart, Tus/*Ter*-induced LTGC is Rad51-independent and is quantitatively increased in the absence of BRCA1, suggesting a non-classical mechanism of initiation (Willis et al., 2014).

Non-homologous TDs of ~2-6 kb—scored as GFP^−^RFP^+^ products (**Figure 1B**)—can be observed during the stalled fork response but not during conventional DSB repair (Willis et al., 2017). Tus/*Ter*-induced TD formation is suppressed by BRCA1 and its binding partners BARD1 and CtIP, but is independent of BRCA2 or Rad51. Remarkably, the genomes of human breast and ovarian cancers lacking *BRCA1*, but not those lacking *BRCA2*, contain abundant small ~10 kb non-homologous TDs (termed “Group 1” TDs), which drive *BRCA1*-linked cancer primarily by disrupting tumor suppressor genes (Menghi et al., 2018; Menghi et al., 2016; Nik-Zainal et al., 2016). Thus, the *Tus/Ter* system recapitulates the phenomenon of Group 1 TD formation in *BRCA1*-linked cancer. Tus/*Ter*-induced TDs arise by an aberrant replication fork restart/replication bypass mechanism and are completed by an end joining step (Willis et al., 2017). Notably, although depletion of either FANCM or BLM has little impact on TD frequencies in wild type cells, depletion of either protein in *Brca1* mutants boosts Tus/*Ter-induced* TDs >10-fold (Willis et al., 2017).

Precisely how FANCM, BLM and BRCA1 interact in stalled fork repair, including in the suppression of Group 1 TDs, remains to be defined. Are the distinct stalled fork repair outcomes noted above reflections of the function or dysfunction of a single overarching activity, or are they independently regulated? In this study, we answer this question and reveal a key role for FANCM in repair pathway choice at stalled forks. We also identify unexpected synthetic lethal interactions between loss-of-function mutations in *Brca1* and *Fancm*.

## Results

### FANCM is recruited to Tus/*Ter*-stalled mammalian replication forks

To study the role of FANCM in stalled fork repair, we engineered mutations of endogenous *Fancm* in a mouse embryonic stem (ES) cell line that carries a conditional allele of *Brca1* (Willis et al., 2014; Willis et al., 2017; Xu et al., 1999b) (discussed below) and contains a single copy of a 6x*Ter*-HR reporter targeted to one allele of *Rosa26* on chromosome 6 (**Figure 1B**) (Willis et al., 2014; Willis et al., 2017). In an effort to generate *Fancm* derivatives of these cells, we used CRISPR/Cas9 with dual sgRNA incisions to engineer an 85 bp deletion within exon 2, introducing a frame-shift early in the *Fancm* open reading frame (ORF). We derived multiple independent clones with potentially biallelic *Fancm*^Δ85^ mutations, as well as multiple independent parallel Cas9/sgRNA-exposed clones that retained wild type *Fancm*—(here termed *Fancm*^+/+^ cells; **Figures 1C**, **S1A** and **S1B**).

CRISPR/Cas9-engineered “*Fancm*^Δ85/Δ85^” clones might contain one precise 85 bp deletion and one larger deletion affecting the second *Fancm* allele (Kosicki et al., 2018). We therefore adopt the term *Fαncm*^Δ85/Δ^ to describe these clones. We used whole genome sequencing to determine the genotype of *Fancm*^Δ85/Δ^ *Ter*-HR reporter clone #39, comparing it to *Fancm*^+/+^ clone #48 (**Figure S1C**). In addition to the expected *Brca1* conditional genotype (described later), we observed heterozygous *Fancm* deletions—the planned 85bp deletion and a larger deletion of 2544 bp sharing its 5’ edge with the Δ85 allele but extending into the 2^nd^ intron. This deletion is expected to disrupt the *Fancm* ORF as effectively as the planned Δ85 bp exon 2 frame-shift. Three independent *Fancm*^Δ85/Δ^ clones revealed greatly reduced *Fancm* mRNA levels compared to *Fancm*^+/+^ cells, suggestive of nonsense-mediated decay (**Figure 1D**). Full-length FANCM protein was undetectable in all three *Fancm*^Δ85/Δ^ clones (**Figures 1E**). To test for sensitivity of *Fancm*^Δ85/Δ^ cells to the ICL-inducing agent mitomycin C (MMC), we exposed a mixed culture of ~90% unmarked (GFP^−^) *Fancm*^Δ85/Δ^ clone 39 cells and ~10% GFP^+^-marked wild type cells to titrated doses of MMC for 72 hours (**Figure 1F**). GFP^+^ cells were progressively enriched at higher doses of MMC, indicating a relative fitness advantage of the GFP^+^ wild type cells over the GFP^−^ *Fancm*^Δ85/Δ^ cells. No equivalent enrichment was observed when ~10% GFP^+^ wild type cells were mixed with ~90% GFP^−^ *Fancm^+/+^* cells. Thus, *Fancm^Δ85/Δ^* cells are sensitive to MMC, consistent with previous studies of *FANCM* (Deans and West, 2009; Rosado et al., 2009). Taken together, these findings suggest that *Fancm*^Δ85/Δ^ is null for *Fancm*.

FANCM localizes to stalled replication forks, but its precise distribution in living in mammals has not previously been measurable. We used chromatin-immunoprecipitation (ChIP) to determine whether FANCM accumulates at the Tus/*Ter* RFB. To this end, we generated *fancum*^Δ85/Δ^ and control Cas9/sgRNA treated wild type derivatives of conditional *Brca1* ES cells that contains a *Rosa26*-targeted *GFP* allele with embedded *Ter* and *I-SceI* sites but lacks the second *GFP* repeat or synthetic *RFP* exons (Willis et al., 2018). We observed strong enrichment of FANCM at the 6x*Ter* array in Tus-transfected *Fancm*^+/+^ cells compared to parallel cultures that received empty vector (**Figure 1G**). In contrast, Tus-transfected *Fancm*^Δ85/Δ^ cells revealed no FANCM-specific ChIP signal. We also noted recruitment of FANCM to chromatin near an I-SceI-induced DSB (**Figure 1H**). Interestingly, the FANCM signals at both a *Tus/Ter* RFB and an I-SceI-induced DSB were restricted to <1 kb either side of the inducing lesion. Thus, FANCM localizes specifically to stalled fork structures at a Tus/*Ter* RFB and in close proximity to an I-SceI-induced DSB.

### FANCM regulates three distinct pathways of stalled fork repair

To study the impact of *Fancm* loss on Tus/*Ter*-induced repair, we measured repair in parallel cultures of 3 independent *Fancm*^Δ85/Δ^ clones (#39, 44 and 49) and 3 independent Fancm^+/+^ clones (#46, 47 and 48) of *Ter*-HR reporter cells. We transfected cells, in parallel, with Tus or I-SceI-expression vectors or empty vector (EV), and measured repair products 72 hours later (see STAR methods). Notably, all *Fancm*^Δ85/Δ^ clones revealed a ~4-fold reduction in Tus/*Ter*-induced STGC and a ~2-fold increase in LTGC, compared to Fancm^+/+^ controls; indeed, loss of *Fancm* dramatically skewed the proportion of LTGC:Total HR products at Tus/*Ter* from ~10% to ~70% (**Figure 2A**). Neither Fancm^+/+^ nor *Fancm*^Δ85/Δ^ cells revealed Tus/*Ter*-induced TDs (GFP^−^RFP^+^ products) in the absence of BRCA1 depletion. *Fancm*^Δ85/Δ^ cells revealed a modest reduction in I-SceI-induced STGC, but no change in I-SceI-induced LTGC (**Figure 2B**). Thus, *Fancm*^Δ85/Δ^ cells reveal HR defects that are specific to stalled fork repair, as revealed by severe impairment of STGC and elevated LTGC frequencies.

**Figure 2.**
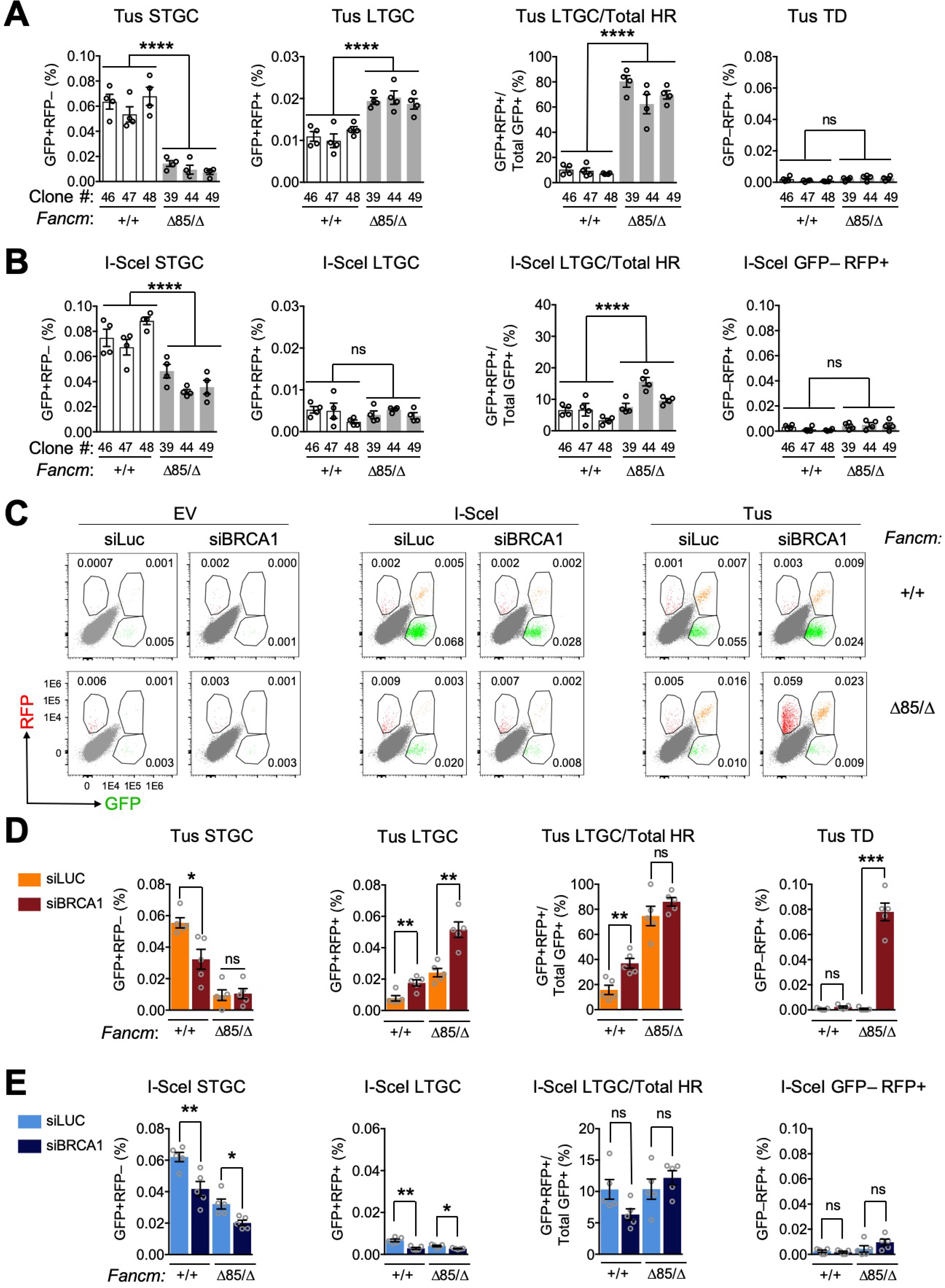
FANCM regulates three distinct pathways of stalled fork repair. See also Figure S2. **A.** Tus/*Ter*-induced HR in three *Fancm^+/+^* (white bars) clones *vs*. three *Fancm*^Δ85/Δ^ (gray bars) clones. Data shows mean of four independent experiments (n=4). Error bars: standard error of the mean (s.e.m.). ****: P < 0.0001 by one-way ANOVA. ns: not significant. **B.** I-SceI-induced repair measured in parallel in same experiment as Panel A. **C.** Representative raw FACS data (uncorrected for transfection efficiency) for *Fancm^+/+^* and *Fancm*^Δ85/Δ^ 6x*Ter*-HR reporter clones co-transfected with either empty vector (EV), I-SceI or Tus expression vectors and siRNAs as shown. FACS plots produced from pooled technical duplicates, n=4. Numbers indicate percentages. **D.** Tus/*Ter*-induced repair in *Fancm^+/+^* clone #48 and *Fancm*^Δ85/Δ85^ clone #39 co-transfected with Tus expression plasmid and siRNAs as shown. Data shows mean values, n=4. Error bars: s.e.m. *: P < 0.05, **: P < 0.01, ***: P < 0.001 by Student’s *t*-test (see STAR Methods). **E.** I-SceI-induced repair measured in parallel in same experiment as Panel D.

To study the relationship between *Fancm* and BRCA1, we used siRNA to deplete BRCA1 in *Fancm*^Δ85/Δ^ or control *Fancm^+/+^ Ter*-HR reporter cells co-transfected with Tus, I-SceI or control empty vectors (see STAR methods and **Figure S2A**). We used transfection of siRNA to luciferase (siLUC) in parallel samples as a mock-depletion control for siBRCA1. **Figure 2C** shows representative raw data, uncorrected for transfection efficiency, from these experiments. Depletion of BRCA1 in Fancm^+/+^ *Ter*-HR reporter cells produced the expected effects of reduced Tus/*Ter*-induced STGC and increased LTGC (**Figure 2D**) (Willis et al., 2014). BRCA1 depletion did not further reduce Tus/*Ter*-induced STGC in *Fancm*^Δ85/Δ^ cells, but it did increase Tus/*Ter*-induced LTGC. *Fancm*^Δ85/Δ^ cells depleted of BRCA1 revealed strong induction of Tus/*Ter*-induced TDs (**Figure 2D**), recapitulating the previously noted synergistic relationship between FANCM loss and BRCA1 loss in TD formation (Willis et al., 2017). Defects in I-SceI-induced STGC caused by loss of *Fancm* and BRCA1 depletion appeared to be additive, suggesting possible differences in the relationships between FANCM and BRCA1 in STGC regulation at stalled forks and at DSBs (**Figure 2E**). We noted no significant induction of I-SceI-induced GFP^−^RFP^+^ products in *Fancm*^Δ85/Δ^ cells depleted of BRCA1, confirming specificity of TD induction for the stalled fork response. Indeed, our unpublished work shows that I-SceI-induced GFP^−^RFP^+^ products are not TDs (data not shown).

### *Fancm* hemizygous cells retain wild type stalled fork repair phenotypes

To study the contribution of individual FANCM domains to stalled fork repair, we engineered in-frame mutations in the endogenous *Fancm* gene in *Brca1* conditional *Ter*-HR reporter cells. This method has the advantage of preserving physiological regulation of gene expression. To avoid confounding effects of potentially heterozygous Cas9-induced mutations affecting the two *Fancm* alleles, we first generated *Brca1* conditional *Ter*-HR reporter cells that are strictly hemizygous for *Fancm*. We used CRISPR/Cas9 with dual sgRNA targeting sites within exons 2 and 23 of *Fancm* to delete a 53.2 kb fragment encompassing one entire *Fancm* allele (here termed *Fancmr*^−^; **Figures S2B** and **S2C**). Sequencing of a selected *FancnΔ* clone revealed no Cas9-induced indels at the sgRNA target sites of the residual wild type *Fancm* allele, and *Fancm* expression matched that of *Fancm*^+/+^ cells, implying transcriptional compensation for hemizygosity (**Figures S2C** and **S2D**). The *Fancm*^+/−^ *Brca1* conditional *Ter*-HR reporter clone revealed wild type repair frequencies; *Fancm*^Δ85/Δ^ cells served as *Fancm*-defective controls in these experiments (**Figure S2E**).

### The FANCM-FA core complex interaction mediates Tus/*Ter*-induced STGC

The FANCM MM1 domain interacts with FANCF and is implicated in recruitment of the FA core complex to stalled forks (Deans and West, 2009). We used Cas9-dual sgRNA incisions to generate an in-frame 366 bp deletion of the MM1 coding region of exon 14 in the residual wild type *Fancm* allele of *Fancm*^+/−^ *Ter*-HR reporter cells (**Figure 3A** and **Figure S3A**). We studied four independent clones of each genotype of *Fancm*^ΔMM1/−^ (#13, 19, 47 and 77) and equivalently Cas9/sgRNA-exposed *Fancm*^+/−^ cells (#01, 07, 14 and 16); each clone’s genotype was confirmed by sequencing. In *Fancm*^ΔMM1/−^ clones, MM1-encoding mRNA was undetectable, whereas MM2-encoding mRNA was detected at normal levels, as was the FANCM protein (**Figures 3B** and **3C**). Consistent with a previous study, *Fancm*^ΔMM1/−^ cells revealed increased sensitivity to MMC (Deans and West, 2009) (**Figure 3D**). We used ChIP to study the recruitment of FA core complex components to Tus/*Ter* in *Fancm*^ΔMM1/−^ and *Fancm*^+/−^ cells that contain a 6x*Ter*-I-SceI cassette at *Rosa26* that lacks a *GFP* repeat (Willis et al., 2018). We detected robust Tus-dependent recruitment of FANCA and FANCL to Tus/*Ter* in *Fancm*^+/−^ cells, but no recruitment in *Fancm*^ΔMM1/−^ cells (**Figure 3E**). This shows that the FANCM MM1 domain is required for stable interaction of the FA core complex with Tus/*Ter*-stalled forks.

**Figure 3.**
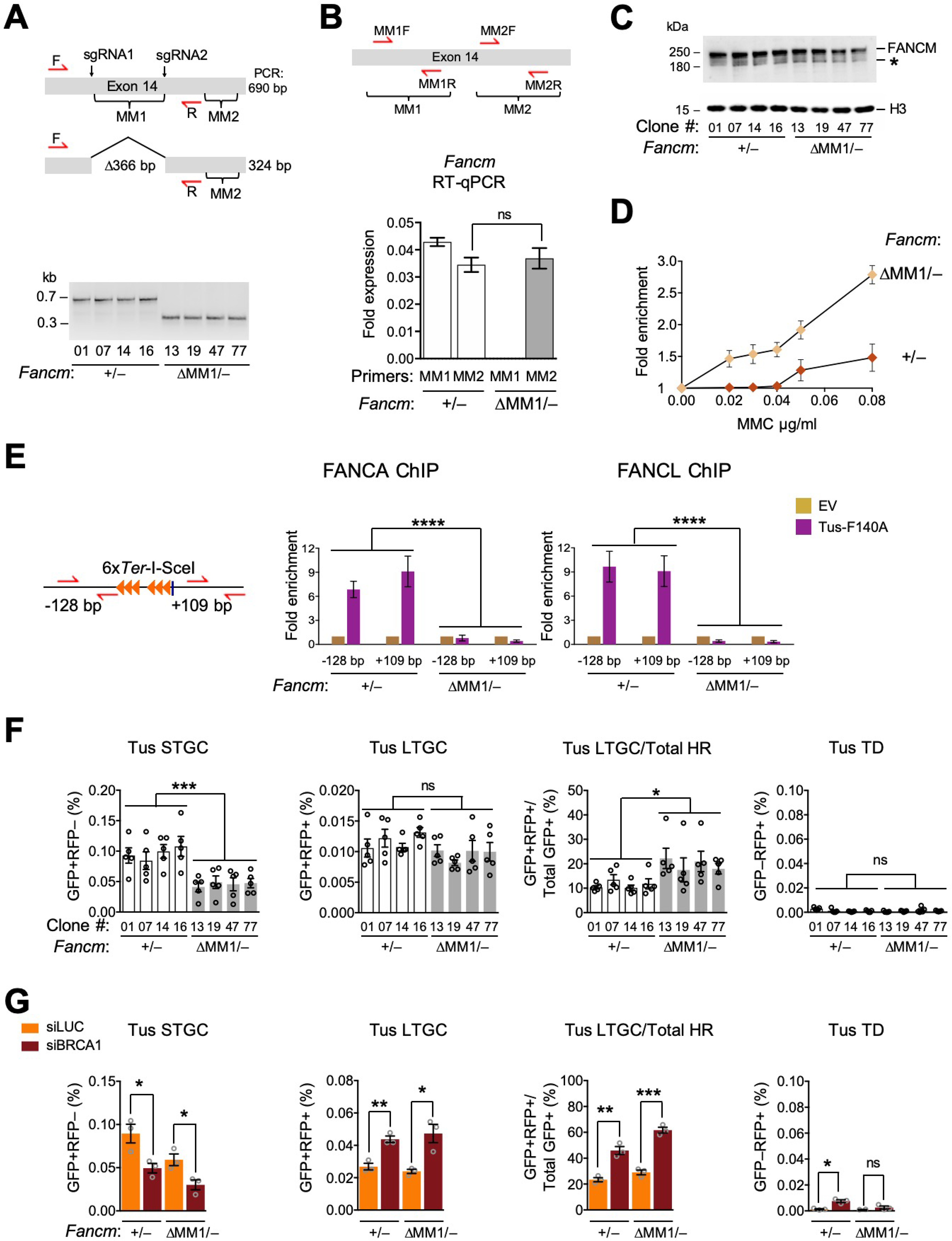
The FANCM-FA core complex interaction mediates Tus/*Ter*-induced STGC. See also Figure S3. **A.** Plan for generating 336 bp in-frame deleted *Fancm*^ΔMM1^ allele. Red half-arrow heads: genotyping primers as shown. Gel: PCR products of *Fancm*^+/−^ and *Fancm*^ΔMM1/−^ clones. **B.** RT qPCR analysis of MM1 and MM2 encoding mRNA in *Fancm*^+/−^ and *Fancm*^ΔMM1/−^ clones. Cartoon shows PCR primer pairs. Data normalized to *Gapdh* mRNA using the 2^−ΔCT^ method (n=3) and analyzed by Student’s *t*-test. Error bars: standard deviation. ns: not significant. **C.** Chromatin-bound FANCM abundance in four *Fancm*^+/−^ and four *Fancm*^ΔMM1/−^ clones. *: nonspecific band. H3: Histone H3 loading control. **D.** Proliferative competition assay in presence of Mitomycin C (MMC), measuring enrichment of GFP^+^ *Fancm vs*. GFP^−^ *Fancm*^ΔMM1/−^ cells. Data, normalized to 0 μg/mL MMC, shows mean value (n=3). Error bars: standard deviation. **E.** ChIP analysis of FANCA and FANCL at Tus/*Ter* RFB in *Fancm*^+/−^ and *Fancm*^ΔMM1/−^ cells (n=3). Elements and data analysis as in Figure 1G. ****: P < 0.0001 by one-way ANOVA. **F.** Tus/*Ter*-induced HR in four *Fancm*^+/−^ (white bars) clones *vs*. four *Fancm*^ΔMM1/−^ (gray bars) clones. Data shows mean values (n=5). Error bars: s.e.m. *: P < 0.05, ***: P < 0.001 by one-way ANOVA. ns: not significant. **G.** Tus/*Ter*-induced repair in *Fancm vs*. *Fancm*^ΔMM1/−^ clones co-transfected with Tus expression plasmid and siRNAs as shown. Data shows mean values, n=3. Error bars: s.e.m. *: P < 0.05, **: P < 0.01, ***: P < 0.001 by Student’s *t*-test.

We analyzed stalled fork and DSB repair functions of each of the above-noted *Fancm*^ΔMM1/−^ and isogenic *FancnΔ* clones. Tus/*Ter*-induced STGC was reduced in *Fancm*^ΔMM1/−^ cells compared to controls (**Figure 3F**). However, this defect was less severe that the defect in Tus/*Ter*-induced STGC noted in *Fancm*^Δ85/Δ85^ cells (compare with **Figure 2A**). Tus/*Ter*-induced LTGC was unaltered by deletion of MM1, producing a compensatory increase in the LTGC/Total HR ratio. No TD induction was noted. Depletion of BRCA1 in *Fancm*^ΔMM1/−^ cells exacerbated the defect in Tus/*Ter*-induced STGC and elevated LTGC (**Figure 3G**). Therefore, BRCA1 controls Tus/*Ter*-induced HR independently of the FANCM MM1 domain. In contrast to *Fancm*^Δ85/Δ^ cells, *Fancm*^ΔMM1/−^ cells revealed no induction of Tus/*Ter*-induced TDs following BRCA1 depletion (**Figure 3G**). We noted modest, proportionate reductions in I-SceI-induced STGC and LTGC in *Fancm*^ΔMM1/−^ cells, but no other repair defects (**Figure S3B**). BRCA1 depletion diminished I-SceI-induced STGC and LTGC proportionately in *Fancm*^ΔMM1/−^ *Fancm*^ΔMM1/−^ cells. Taken together, these results show that the FANCM MM1 domain positively regulates STGC at stalled forks, but additional FANCM elements also contribute to this function. The lack of LTGC dysregulation in *Fancm*^ΔMM1/−^ cells and the absence of Tus/*Ter*-induced TDs following BRCA1-depletion identify *Fancnm^ΔMM1^* as a separation-of-function allele that discriminates control of STGC from other FANCM-mediated stalled fork repair functions.

### The FANCM-BLM interaction suppresses LTGC and TD formation at stalled forks

The FANCM MM2 domain interacts with the BTR complex *via* direct interactions with RMI1 (**Figure 1A**) (Deans and West, 2009). We used Cas9-dual sgRNA incisions to generate an inframe 114 bp deletion of the MM2-encoding region of exon 14 in the residual wild type *Fancm* allele of *Fancm*^+/−^ *Ter*-HR reporter cells (**Figure 4A** and **Figure S4A**). We studied three independent clones of each genotype of *Fancm*^ΔMM2/−^ (#03, 06 and 09) and equivalently Cas9/sgRNA-exposed *FancnΓ* cells (#22, 26, 27); each clone’s genotype was confirmed by sequencing. In *Fancm*^ΔMM2/−^ clones, MM2-encoding mRNA was undetectable, whereas MM1-encoding mRNA was detected at normal levels, as was the FANCM protein (**Figures 4B** and **4C**). *Fancm*^ΔMM2/−^ cells revealed only modest sensitivity to MMC (**Figure 4D**). We used ChIP to study the recruitment of BLM to Tus/*Ter* in *Fancm*^ΔMM2/−^ and control *Fancm* cells that contain a minimal 6x*Ter*-I-SceI cassette at *Rosa26* (Willis et al., 2018). We detected robust Tus-dependent recruitment of BLM to Tus/*Ter* in *Fancm^+/+^* and *Fancm*^+/−^ cells, but no recruitment in *Fancm*^Δ85/Δ^ cells (**Figure 4E**). The BLM ChIP signal in *Fancm*^ΔMM2/−^ cells was reduced but not absent. This shows that *Fancm* is required for BLM recruitment to a Tus/*Ter* RFB, and the MM2 domain mediates this function. However, additional FANCM elements also contribute to BLM recruitment.

**Figure 4.**
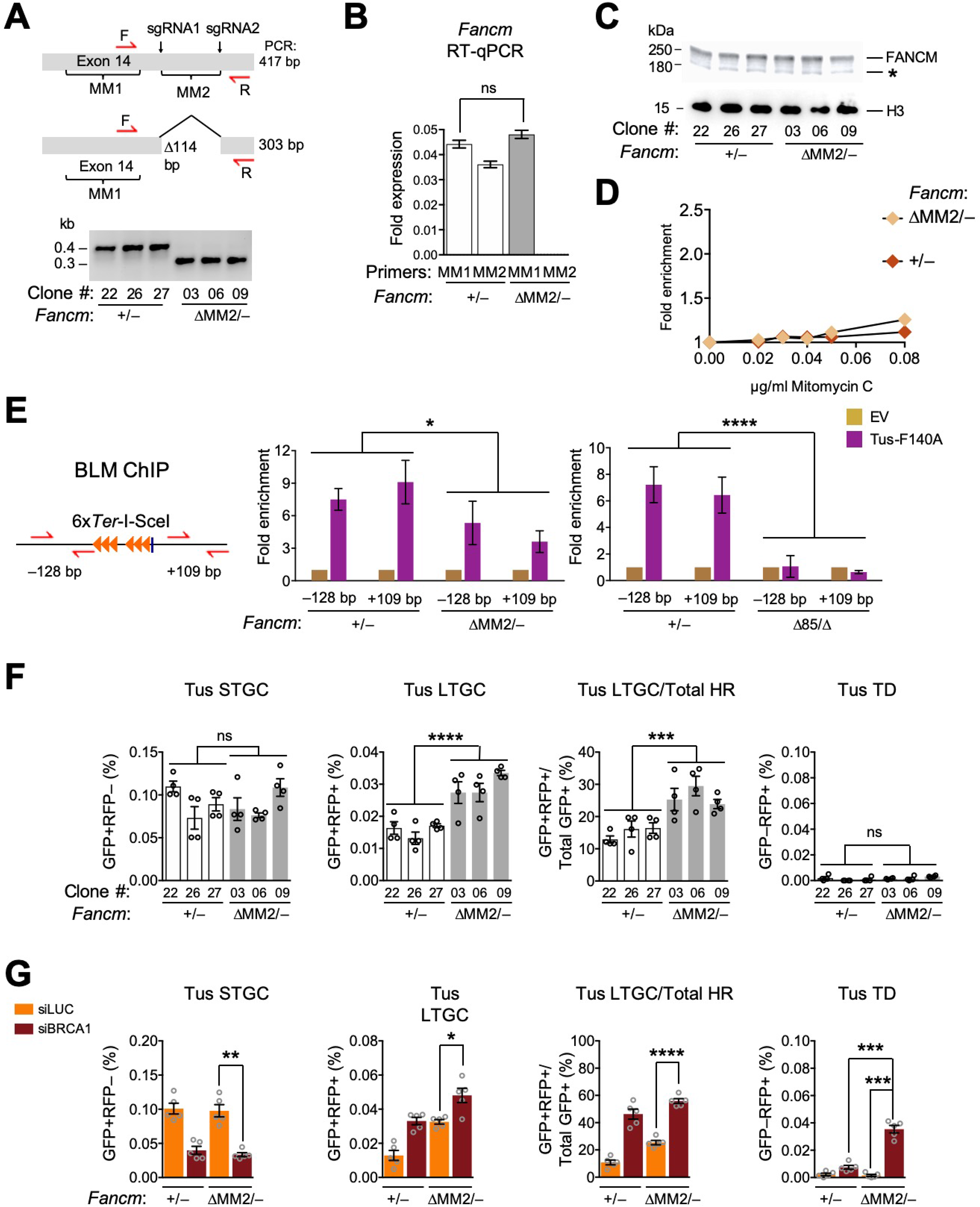
The FANCM-BLM interaction suppresses LTGC and TD formation at stalled forks. See also Figure S4. **A.** Plan for generating 336 bp in-frame deleted *Fancm*^ΔMM2^ allele. Red half-arrow heads: genotyping primers as shown. Gel: PCR products of *Fancm*^+/−^ and *Fancm*^ΔMM1/−^ clones. **B.** RT qPCR analysis of MM1 and MM2 encoding mRNA in *Fancm*^+/−^ and *Fancm*^ΔMM2/−^clones. Data shows mean values (n=3) normalized to *Gapdh* mRNA using the 2^−ΔCT^ method. Analysis by Student’s *t*-test. Error bars: standard deviation. ns: not significant. **C.** Abundance of chromatin-bound FANCM in three *Fancm*^+/−^ and three *Fancm*^ΔMM2/−^ clones. H3: histone H3 loading control. **D.** Proliferative competition assay in presence of Mitomycin C (MMC), measuring enrichment of GFP^+^ *Fancm*^+/−^ *vs*. GFP^−^ *Fancm*^ΔMM2/−^ cells. Data, normalized to 0 μg/mL MMC, shows mean value (n=3). Error bars (standard deviation) are obscured by data points. **E.** ChIP analysis of BLM at Tus/*Ter* RFB in *Fancm*^+/−^ and *Fancm*^ΔMM2/−^ cells (n=3). Elements and data analysis as in Figure 1G. *: P < 0.05, ****: P < 0.0001 by one-way ANOVA. **F.** Tus/*Ter*-induced HR in three *Fancm*^+/−^ (white bars) clones *vs*. four *Fancm*^ΔMM2/−^ (gray bars) clones. Data shows mean values (n=4). Error bars: s.e.m. ***: P < 0.001, ****: P < 0.0001 by one-way ANOVA. ns: not significant. **G.** Tus/*Ter*-induced repair in *Fancm*^+/−^ *vs. Fancm*^ΔMM2/−^ clones co-transfected with Tus expression plasmid and siRNAs as shown. Data shows mean values, n=5. Error bars: s.e.m. *: P < 0.05, **: P < 0.01, ***: P < 0.001, ****: P < 0.0001by Student’s *t*-test.

We studied stalled fork and DSB repair functions of each of the above-noted *Fancm*^ΔMM2/−^ and isogenic *Fancm* clones. *Fancm*^ΔMM2/−^ cells revealed unaltered Tus/*Ter*-induced STGC compared to *Fancn*^+/−^ controls and no Tus/*Ter*-induced TDs in the absence of BRCA1 depletion; however, Tus/*Ter*-induced LTGC was elevated ~1.5-fold (**Figure 4F**). Following depletion of BRCA1, we noted reduction in Tus/*Ter*-induced STGC and further increases in LTGC in both *Fancm*^+/−^ and *Fancm*^ΔMM2/−^ cells, indicating that BRCA1 functions independently of the FANCM MM2 domain in stalled fork HR. Notably, BRCA1-depletion boosted Tus/*Ter*-induced TDs in *Fancm*^ΔMM2/−^ cells in comparison to *Fancm*^+/−^ controls, albeit to frequencies lower than those observed in BRCA1-depleted *Fancm*^Δ85/Δ^ cells (**Figure 4G;** compare with **Figure 2D**). No significant alterations in I-SceI-induced HR were noted in *Fancm*^ΔMM2/−^ cells (**Figures S4B** and **S4C**). These findings indicate that MM2 specifically suppresses Tus/*Ter*-induced LTGC and TD formation (the latter in cells lacking BRCA1, but has no impact on STGC at stalled forks. *Fancm*^Δ*MM2*^ is therefore a separation-of-function allele that discriminates suppression of LTGC and TD formation from STGC control in stalled fork repair.

### FANCM ATP hydrolysis mutant is defective for FANCM-mediated stalled fork repair

To determine the contribution of FANCM ATPase function to stalled fork repair, we used dual Cas9/sgRNA incisions to engineer an in-frame 66 bp deletion within the FANCM ATPase domain-encoding region of exon 2 of *Fancm*^+/−^ *Ter*-HR reporter cells (**Figure 5A** and **Figure S5A**). The resulting *Fancm*^ΔDEAH/−^ product lacks 22 amino acid residues spanning the DEAH motif of the Walker B box and is predicted to be defective for ATP hydrolysis. We studied three independent clones of each genotype of *Fancm*^ΔDEAH/−^ (#16, 55 and 67) and equivalently Cas9/sgRNA-exposed *Fancn*^+/−^ cells (#12, 13 and 14); each clone’s genotype was confirmed by sequencing. In *Fancm*^ΔDEAH/−^ clones, mRNA sequence encoding the deleted 22 amino acids spanning the DEAH motif was undetectable, whereas MM2-encoding mRNA sequences were detected at normal levels, as was the FANCM protein (**Figures 5B** and **5C**). We used ChIP to study the recruitment of FANCM and BLM to Tus/*Ter* in *Fancm*^ΔDEAH/−^ and *Fancm*^+/−^ cells that contain a 6x*Ter* and *I-SceI* site at *Rosa26* but lack one of the *GFP* repeats (Willis et al., 2018). In both cell types, we detected robust Tus-dependent recruitment of BLM (**Figure 5D**). Interestingly, the FANCM ΔDEAH mutant protein accumulated to higher levels at Tus/*Ter* than wild type FANCM (**Figure 5D**). Thus, the motor function of FANCM is dispensable for BLM recruitment to Tus/*Ter*-stalled forks, but appears to be required for the timely release of FANCM from the stalled fork.

**Figure 5.**
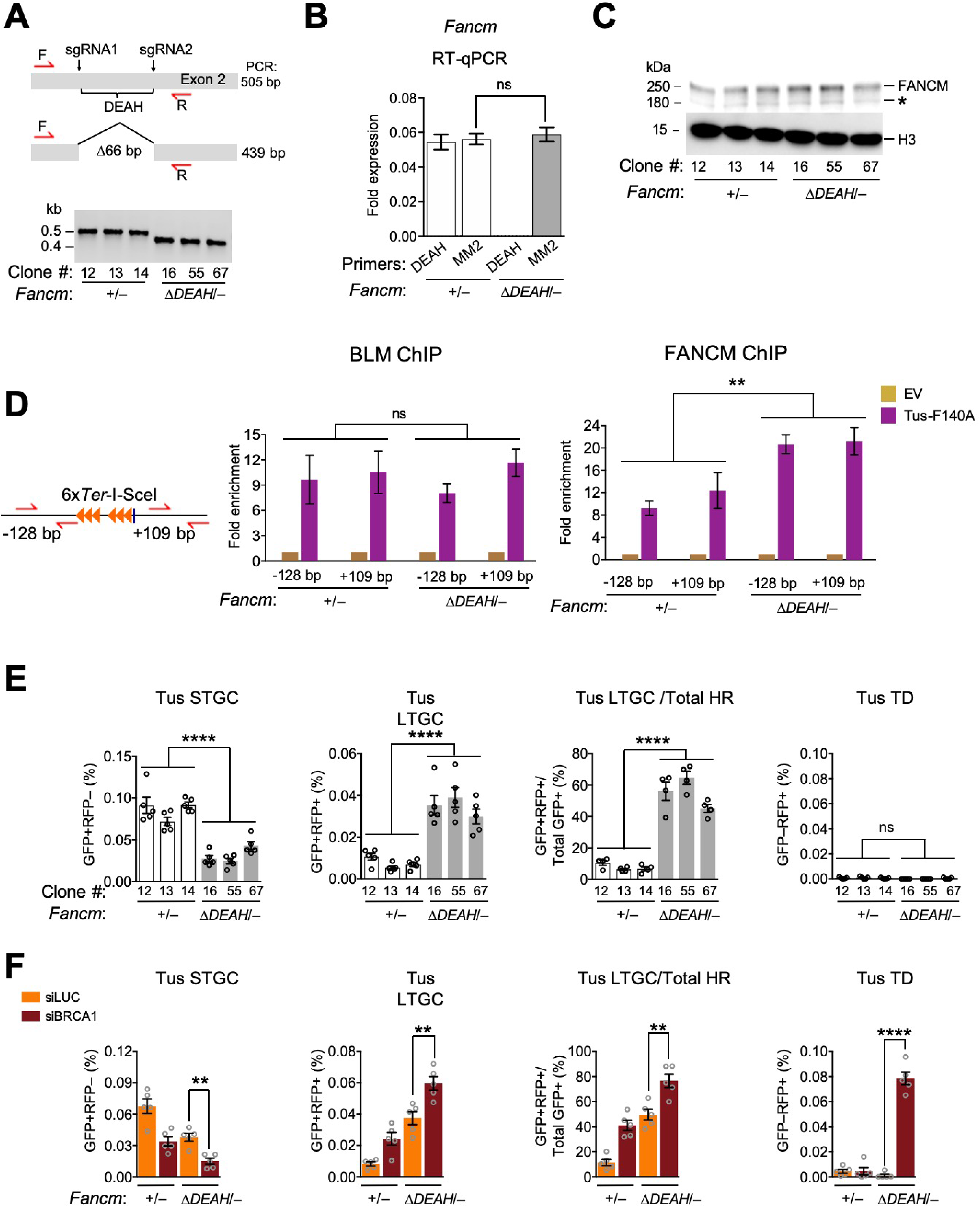
FANCM ATP hydrolysis mutant is defective for FANCM-mediated stalled fork repair. See also Figure S5. **A.** Plan for generating 66 bp in-frame deleted *Fancm*^ΔDEAH^ allele. Red half-arrow heads: genotyping primers as shown. Gel: PCR products of *Fancní* and *Fancm*^ΔDEAH/−^ clones. **B.** RT qPCR analysis of DEAH and MM2 encoding mRNA in *Fancm* and *Fancm*^ΔDEAH/−^clones. Data normalized to *Gapdh* mRNA using the 2^−ΔCT^ method (n=3) and analyzed by Student’s *t*-test. Error bars: standard deviation. ns: not significant. **C.** Abundance of chromatin-bound FANCM in three *Fancm* and three *Fancm*^ΔDEAH/−^ clones. H3: histone H3 loading control. **D.** ChIP analysis of BLM at Tus/*Ter* RFB in *Fancm* and *Fancm*^ΔDEAH/−^ cells (n=3). Elements and data analysis as in Figure 1G. **: P < 0.01, ns: not significant by one-way ANOVA. **E.** Tus/*Ter*-induced HR in three *Fancm*^+/−^ (white bars) clones *vs*. three *Fancm*^ΔDEAH/−^ (gray bars) clones. Data shows mean values (n=5). Error bars: s.e.m. ****: P < 0.0001 by oneway ANOVA. ns: not significant. **F.** Tus/*Ter*-induced repair in *Fancm vs*. *Fancm*^ΔDEAH/−^ clones co-transfected with Tus expression plasmid and siRNAs as shown. Data shows mean values, n=5. Error bars: s.e.m. **: P < 0.01, ****: P < 0.0001by Student’s *t*-test.

We studied stalled fork and DSB repair functions of each of the above-noted *Fancm*^ΔDEAH/−^ and isogenic *Fancm*^+/−^ clones. Notably, *Fancm*^ΔDEAH/−^ clones revealed a ~3-fold reduction in Tus/*Ter*-induced STGC, a ~3-fold increase in LTGC (GFP^+^RFP^+^ products), compared to *Fancn*^+/−^ controls, and a corresponding skewing of the proportion of LTGC:Total HR products at Tus/*Ter* from ~10% in *Fancim*^+/−^ cells to ~50% in *Fancm*^ΔDEAH/−^ cells (**Figure 5E**). *Fancm*^ΔDEAH/−^ cells revealed no Tus/*Ter*-induced TDs in the absence of BRCA1 depletion (**Figure 5E**), and showed a modest reduction in I-SceI-induced STGC, but no change in I-SceI-induced LTGC (**Figure S5B**). BRCA1 depletion exacerbated the defect in Tus/*Ter*-induced STGC and increased LTGC in *Fancm*^ΔDEAH/−^ cells. Thus, *Fancm*^ΔDEAH/−^ cells reveal primary defects in stalled fork repair and DSB repair similar to those observed in *Fancm*^Δ85/Δ^ cells, with the exception that the STGC defect is further diminished by BRCA1 depletion in *Fancm*^ΔDEAH/−^ cells. This difference raises the possibility that a scaffolding function of FANCM contributes to BRCA1-mediated STGC. There was a striking elevation of Tus/*Ter*-induced TDs in *Fancm*^ΔDEAH/−^ cells depleted of BRCA1, reaching frequencies similar to those observed in *Fancm*^Δ85/Δ^ cells (compare **Figures 5F** and **2D**). In response to an I-SceI-induced DSB, we again noted that BRCA1 acts independently of *Fancm* (**Figures S5C**). In summary, all three FANCM-mediated stalled fork repair pathways that our assays measure are profoundly dysregulated if FANCM is unable to hydrolyze ATP. In contrast, certain scaffolding functions of FANCM, such as BLM recruitment to stalled forks and a possible enabling role in BRCA1-mediated STGC at stalled forks, are retained in *Fancm*^ΔDEAH/−^ cells.

### Stalled fork repair functions of BLM are *Fancm*-independent

The Bloom’s helicase appears to be intimately connected to FANCM stalled fork repair functions. First, BLM recruitment to Tus/*Ter* is dependent on *Fancm* (**Figure 4E**). Second, FANCM and BLM are the two major suppressors of TD formation in *Brca1* mutant cells (Willis et al., 2017). Third, the FANCM MM2 domain, which interacts with the BTR complex, contributes to BLM recruitment to Tus/*Ter* and is also required for efficient suppression of Tus/*Ter*-induced LTGC and TD formation (**Figures 4E** and **4G**). In apparent contradiction of these connections, *Fancm*^ΔDEAH/−^ cells support BLM recruitment to stalled forks but show repair defects broadly similar to *Fancm*^Δ85/Δ^ cells. These findings raise the question: to what extent are BLM stalled fork repair functions dependent on *Fancm*? To address this, we depleted BRCA1 and BLM either in parallel or together in *Fancm*^ΔMM2/−^, *Fancm*^Δ85/Δ^ or *Fancm*^ΔDEAH/−^ cells, *vs*. appropriate isogenic controls expressing wild type *Fancm* (**Figure S6)**. We equilibrated siRNA levels in different samples by supplementing with siRNA to Luciferase as appropriate (see STAR methods). We previously found that BLM depletion modestly elevates Tus/*Ter*-induced STGC, increases LTGC and boosts TD formation (the last, in cells lacking BRCA1) (Willis et al., 2017).

In all genetic backgrounds, BLM depletion—either with or without BRCA1 co-depletion— increased Tus/*Ter*-induced STGC and LTGC, although these effects did not reach statistical significance in all samples (**Figure S6**). As expected, BLM and BRCA1 co-depletion triggered Tus/*Ter*-induced TD formation in wild type cells (Willis et al., 2017). However, BLM depletion also significantly increased TD frequencies in BRCA1-depleted *Fancm*^ΔMM2/−^, *Fancm*^Δ85/Δ^ and *Fancm*^ΔDEAH/−^ cells, in comparison to controls that received siBRCA1 + siLuciferase but no siBLM (**Figures S6**). The impact of BLM depletion on TD frequencies was approximately additive on all *Fancm* mutant backgrounds, suggesting that BLM can suppress Tus/Ter-induced LTGC and TD formation independently of *Fancm*. In the same set of experiments, no consistent effects of BLM depletion on I-SceI-induced HR were noted (**Figure S7**). These results suggest that, although the bulk recruitment of BLM to stalled forks requires *Fancm*, BLM can affect stalled fork repair even in presumptively null *Fancm*^Δ85/Δ^ cells, where BLM is not detected at stalled forks by ChIP. These *Fancm*-independent repair functions of BLM may involve transient stalled fork-BLM interactions that are not readily detectable by ChIP.

### Synthetic lethal interaction between *Brca1* and *Fancm* mutations

To study the genetic relationships between loss-of function mutations in *Fancm* and *Brca1*, we sought to conditionally delete wild type *Brca1* in *Fancm* mutant cells. The conditional allele, *Brca1^fl^*, contains *loxP* sites flanking a large in-frame central exon, commonly termed “exon 11” (although it is in fact the 10^th^ exon of *Brca1*) (Miki et al., 1994; Willis et al., 2014; Willis et al., 2017; Xu et al., 1999a). Cre-mediated deletion of exon 11 generates *Brca1^Δ^*, a hypomorphic allele (**Figure 6A**). The second *Brca1* allele, *Brca1^11^*, contains a hygromycin resistance gene in place of the 3’ half of exon 11 that disrupts expression of endogenous *Brca1. Brca1*^fl/11^ ES cells and their Cre-treated *Brca1*^Δ/11^ derivatives grow at equivalent rates *in vitro*, but *Brca1*^Δ/11^ cells reveal HR defects and exhibit synergistic increases in Tus/*Ter*-induced TDs when depleted of either FANCM or BLM (Willis et al., 2014; Willis et al., 2017).

**Figure 6.**
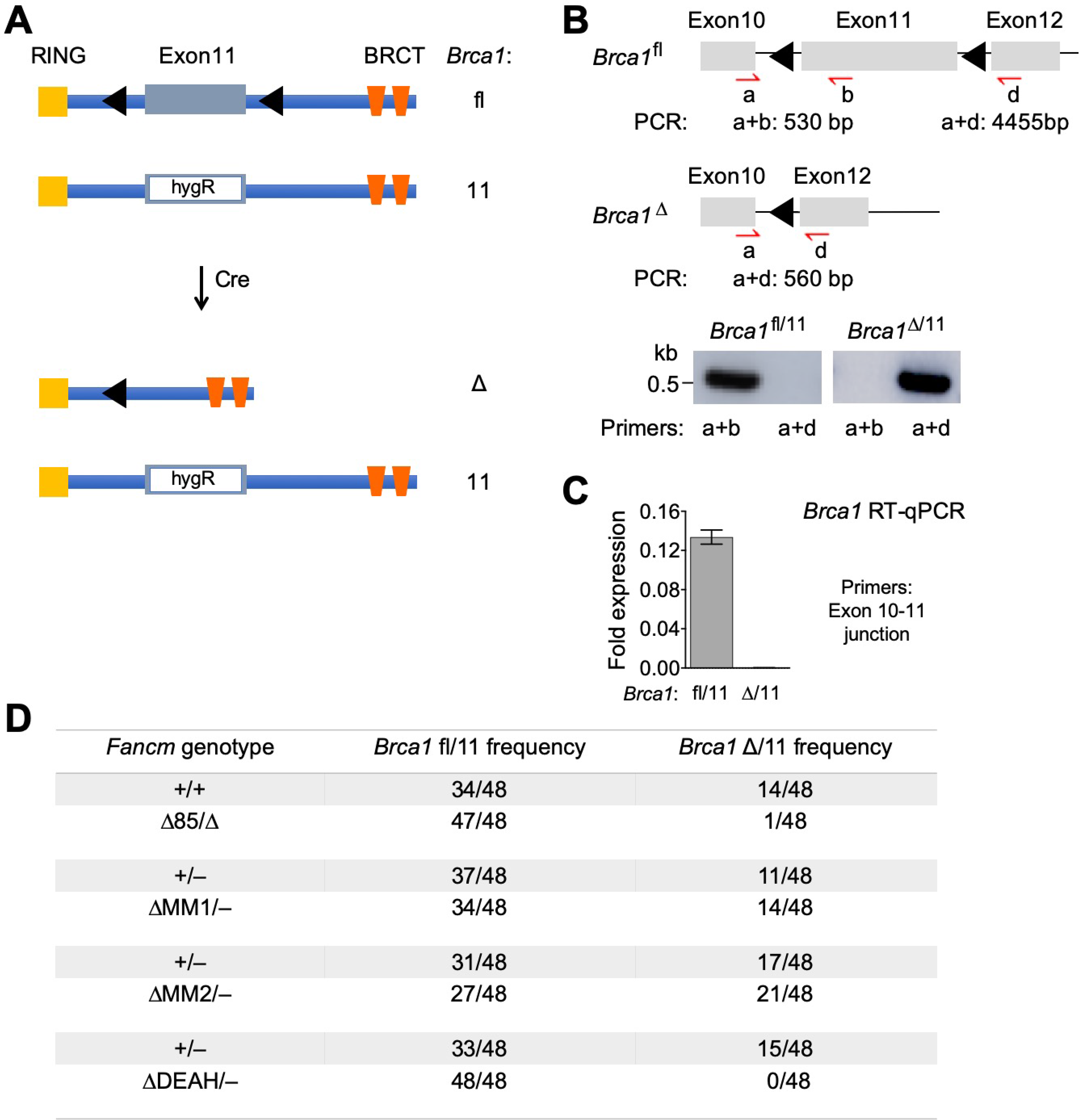
Synthetic lethal interaction between *Brca1* and *Fancm* mutations. See also Figure S8. **A.** BRCA1 structure in *Brca1*^fl/11^ ES cells. Grey box: *Brca1* exon 11. Black triangles: *loxP* elements. *hygR:* hygromycin resistance gene. Transient Cre exposure converts *Brca1^f^* to Exon11-deleted *Brca1^Δ^* allele. **B.** *Brca1* exons 10-12 in *Brca1*^fl/11^ ES cells with PCR primer positions as indicated. Black triangles: *loxP* elements. Red half-arrowheads: PCR primers. Gel: *Brca1*^fl/11^ and *Brca1^Δ/11^* PCR products. **C.** RT qPCR analysis of *Brca1* Exon 11 encoding mRNA in *Brca1*^fl/11^ and *Brca1*^Δ/11^ clones. *Brca1* gene expression level was normalized to *Gapdh* using the 2^−ΔCT^ method (n=3). Error bars: s.e.m. **D.** *Brca1*^fl/11^ and *Brca1*^Δ/11^ allele recovery in unselected clones following transduction of *Brca1*^511^ *Fancm*^Δ85Δ^, *Brca*^fl/11^ *Fancm*^ΔMM15^, *Brca*^fl/11^ *Fancm*^ΔMM25^, or *Brca1*^fl/11^ *Fancm*^ΔDEAH/−^ cells with adenovirus expressing the Cre recombinase.

We exposed cultures of *Brca1*^fl/11^ *Ter*-HR reporter ES cells of different *Fancm* genotype to adenovirus encoding the Cre recombinase and screened clones for loss or retention of wild type *Brca1* (**Figures 6B**, **6C** and **6D**; see also STAR methods). In the first experiment, we retrieved 14/48 (30%) Cre-treated *Fancm^+/+^ Brca1*^Δ/11^ clones, the remainder being of the genotype *Fancm*^+/+^ *Brca1*^fl/11^. In parallel samples of *Fancm*^Δ85/Δ^ *Brca1*^fl/11^ cells exposed to the same titer of adeno-Cre in the same experiment, we retrieved only one *Fancm*^Δ85/Δ^ *Brca1*^Δ/11^ clone out of 48 tested, the remaining 47 clones being *Fancm*^Δ85/Δ^ *Brca1*^fl/11^ (**Figure 6D**). The solitary *Fancm*^Δ85/Δ^ *Brca1*^Δ/11^ clone (#68) was slow-growing compared to its *Fancm*^Δ85/Δ^ *Brca1*^fl/11^ siblings. We used whole genome sequencing to confirm its genotype (**Figures S8A** and **S8B**).

*Fancm*^Δ85/Δ^ *Brca1*^Δn^ clone #68 revealed levels of Tus/*Ter*-induced STGC and LTGC equivalent to an isogenic *Fancm*^Δ85/Δ^ *Brca1*^fl/11^ clone, suggesting that some compensatory changes had occurred in clone #68 (**Figure S8C**). In contrast, clone #68 revealed higher frequencies of Tus/*Ter*-induced TDs than its *Fancm*^Δ85/Δ^ *Brca*^fl/11^ sibling, either in the presence or absence of BRCA1 co-depletion (**Figure S8C**). Clone #68 also revealed lower levels of I-SceI-induced STGC than the isogenic *Fancm*^Δ/85/Δ^ *Brca1*^fl/11^ clone (**Figure S8D**).

We transduced each different *Fancm* genotype of *Brca1*^fl/11^ *Ter*-HR reporter ES cells with adeno-Cre, using appropriate isogenic control clones, and analyzed the resulting colonies. *Brca1*^Δ/11^ clones were recovered at normal frequencies in *Fancm*^ΔMM1/−^ and *Fancm*^ΔMM2/−^ cells (**Figure 6D**). In contrast, we recovered no *Fancm*^ΔDEAH/−^ *Brca1*^Δ/11^ clones, despite retrieving 15/48 (31%) *Brca1*^Δ/11^ clones from parallel control *Fancm*^+/−^ *Brca1*^fl/11^ cultures in the same experiment. Thus, a *Fancm* ATP hydrolysis-defective mutant is synthetic lethal with *Brca1*^Δ/11^ in mouse ES cells.

## Discussion

Work described here defines in detail how FANCM regulates repair of stalled mammalian replication forks and identifies a synthetic lethal interaction between loss-of-function mutations of *Fancm* and *Brca1*. We show that FANCM positively regulates error-free STGC in response to a Tus/*Ter* RFB, while suppressing error-prone replicative responses of LTGC and TD formation. Prior to this study, the regulatory relationships between error-free and error-prone repair at stalled forks were unclear. Our identification of separation-of-function *Fancm* mutants shows that STGC support and LTGC/TD suppression at stalled forks are regulated in a mutually independent fashion, and that FANCM critically determines the balance of flux through these pathways. Underpinning all FANCM-mediated repair is a requirement for its motor protein function, as revealed by the phenotype of the ATPase-defective *Fancm*^ΔDEAH^ mutant. Notably, the FANCM ΔDEAH protein accumulates to unusually high levels at stalled forks, implicating the FANCM motor function in the timely displacement of FANCM from the stalled fork. “Trapping” of the FANCM ATPase-defective mutant may be an important corollary of *Fancm*^ΔDEAH^ mutant repair defects. Finally, we identify synthetic lethal interactions between

*Fancm* null or ATPase mutants and *Brca1* mutation. Collectively, these findings identify the FANCM ATPase as a potentially “druggable” target for therapy of *BRCA1*-linked cancers.

### FANCM in STGC at stalled forks: the Fanconi anemia pathway

The phenotype of *Fancm*^ΔMM1^ mutant cells shows that the interaction of FANCM with the FA core complex specifically supports error-free STGC, while having no role in the regulation of aberrant replicative stalled fork repair responses. These findings draw an explicit connection between Tus/*Ter*-induced STGC and the FA pathway of replication-coupled ICL repair. Notably, the FA pathway is required for cellular viability in the presence of aldehydes, where the dominant fork-stalling lesions may be DNA-protein crosslinks rather than ICLs (Langevin et al., 2011; Rosado et al., 2011). Perhaps Tus/*Ter* will serve as a useful model of how the FA pathway processes such lesions. Similar to the FA pathway of ICL repair, Tus/*Ter*-induced STGC entails conservative two-ended HR, implicating bidirectional fork arrest as an intermediate (Knipscheer et al., 2009; Raschle et al., 2008; Willis et al., 2014; Zhang and Walter, 2014) (**Figure 7A**). The incision step of replication-coupled ICL repair is mediated by the DNA structure-specific endonuclease-binding scaffold SLX4/FANCP (Hodskinson et al., 2014; Klein Douwel et al., 2014). It will be interesting to determine the role of SLX4 and other FA genes in Tus/*Ter*-induced repair.

**Figure 7.**
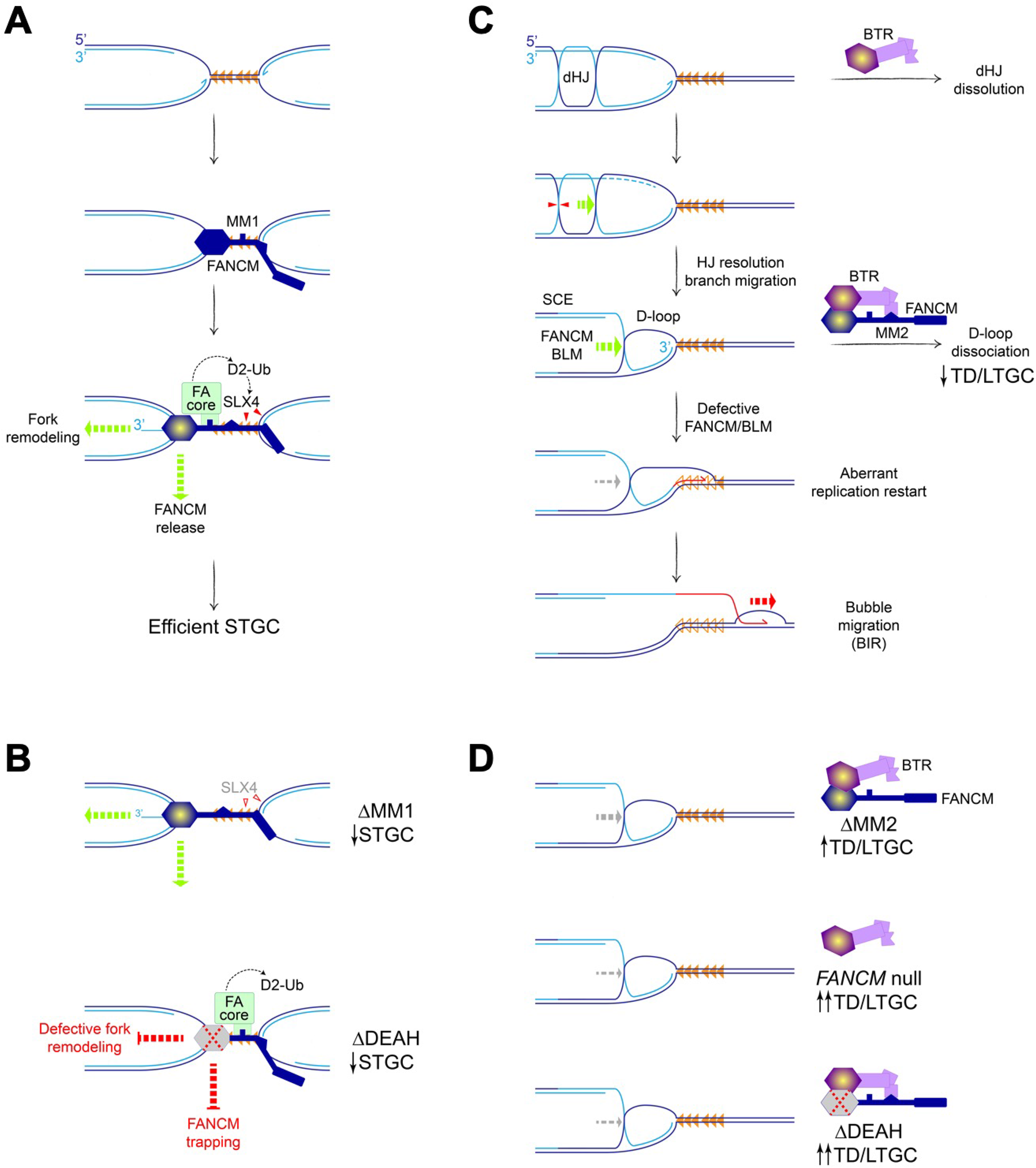
Models of FANCM-mediated stalled fork repair. **A.** FANCM mediates STGC repair of forks bidirectionally arrested at Tus/*Ter* RFB (filled orange triangles). MM1 domain recruits FA core complex, promoting FANCD2 ubiquitination (D2-Ub) and SLX4-mediated incisions (filled red triangles). FANCM motor function (yellow core of hexagon), promotes timely release of FANCM from stalled fork and, possibly, fork remodeling. **B.** Upper cartoon: Defective FA core complex recruitment in *Fancm*^ΔMM1^ mutants suppresses SLX4-mediated incisions (unfilled red triangles) and STGC. Lower cartoon: FANCM trapping and, possibly, defective fork remodeling in *Fancm*^ΔDEAH^ mutants impairs STGC. **C.** FANCM and BLM act in a concerted fashion to suppress aberrant replication fork restart. Hypothetical mechanism of D-loop formation at stalled fork in the absence of a strand exchange step. BLM-TOP3A-RMI1-RMI2 (BTR) complex can dissolve post-replicative double Holliday junction (dHJ). Alternative processing by HJ resolution could generate D-loop at stalled fork, with accompanying sister chromatid exchange (SCE). Red triangles: incisions of HJ resolution. Dashed light blue lines: resected nascent lagging strands at stalled fork. FANCM/BLM-mediated branch migration (green dashed arrow) normally displaces stalled leading nascent strand and dissociates D-loop. Defects in FANCM/BLM allow D-loop to persist, favoring resumption of nascent leading strand synthesis (red half-arrow) with displacement of Tus from *Ter* (empty orange triangles). Engagement of BIR helicase (red dashed arrow) stabilizes aberrant replication restart by BIR/bubble migration. **D.** Impacts of FANCM/BLM dysfunction on D-loop dissociation. Discoordination of FANCM-BTR complex organization in *Fancm*^ΔMM2^ mutants (upper cartoon) diminishes D-loop dissociation/branch migration function, leading to increased TD formation and LTGC. In absence of FANCM (middle panel), or in *Fancm*^ΔDEAH^ mutants (lower panel), this function is more severely impaired, leading to further increases in TD frequencies.

In addition to the MM1 domain, we find that ATP hydrolysis by FANCM is required for efficient Tus/*Ter*-induced STGC. Two distinct mechanisms might explain the STGC defect of *Fancm*^ΔDEAH^ mutants (**Figures 7A** and **7B**). First, “trapping” of an ATPase-defective FANCM mutant at the site of stalled fork processing might interfere with the STGC mechanism. Second, the ability of FANCM to reverse stalled fork model substrates *in vitro* suggests that FANCM may support Tus/*Ter*-induced STGC *via* a fork remodeling step (Gari et al., 2008a; Gari et al., 2008b). Interestingly, an asymmetric fork reversal step is known to accompany FA pathway-mediated ICL repair (Amunugama et al., 2018).

We find that BLM suppresses Tus/*Ter*-induced STGC and, thus, opposes FANCM in this function. Further, the FANCM ΔMM2 mutant, which is impaired in the ability to interact with BLM, exhibits no defects in STGC. These observations suggest that BLM has no direct role in STGC at stalled forks and that FANCM regulates STGC independently of BLM (**Figure 7A**).

### FANCM in LTGC and TD suppression: aberrant replication restart

*Fancm*^ΔMM2^ mutants reveal increased Tus/*Ter*-induced LTGC and TD formation (the latter in the absence of BRCA1). Thus, in contrast to its mechanism of action in STGC, FANCM must interact with the BTR complex for optimal suppression of LTGC and TDs. The mild MMC sensitivity of *Fancm*^ΔMM2/−^ cells suggests that the primary impact of dysregulated LTGC and TD suppression in mouse ES cells may be on mutagenesis/structural variation, rather than on cell viability in response to MMC. In contrast, in HEK 293 cells, the FANCM ΔMM2 mutant revealed MMC sensitivity similar to that of the ΔMM1 mutant, suggesting that the impact of MM2 on MMC sensitivity varies between different cell types (Deans and West, 2009).

In our ChIP studies, we found that the bulk recruitment of BLM to the Tus/*Ter* RFB is dependent on *Fancm*. However, even in *Fancm*^Δ85/Δ^ cells, which are likely *Fancm* null and reveal no BLM ChIP signal at Tus/*Ter*, BLM depletion nonetheless influenced stalled fork repair, increasing the frequencies of LTGC and TD formation in BRCA1-depleted cells. The simplest explanation of this apparent contradiction is that BLM can access stalled fork intermediates in *Fancm*^Δ85/Δ^ cells, but is present at levels below the limits of detection by ChIP. A caveat in our experiments on BLM is the reliance on siRNA-mediated depletion. Greater clarity regarding BLM’s role must await the development of a dedicated genetic system for studying *Blm* in stalled fork repair.

In the experiments reported here, the ability of a *Fancm* mutant to suppress Tus/*Ter*-induced LTGC predicted its ability to suppress TDs. This suggests that Tus/*Ter*-induced LTGC and TD formation—both Rad51-independent processes—may arise by a common mechanism. Tus/*Ter*-induced TDs are products of aberrant replication fork restart (Willis et al., 2014; Willis et al., 2017). Interestingly, replication fork restart in *S. pombe* occurs by a BIR mechanism (Jalan et al., 2019; Nguyen et al., 2015). Taken together, these observations suggest that FANCM/BLM may suppress Tus/*Ter*-induced LTGC and TD formation by disallowing BIR-type replication restart at Tus/*Ter*. A critical step in this restart mechanism would be formation of a stable D-loop at the site of stalling. Given the Rad51-independence of the restart mechanism, it will clearly be relevant to determine the role of *RAD52* in this process. However, in principle, a D-loop could be established at a stalled fork without the need for a strand exchange or annealing step, through the processing of post-replicative Holliday junctions in the vicinity of the fork (**Figure 7C**). Both FANCM and BLM mediate branch migration of Holliday junctions *in vitro*, and the BTR complex additionally mediates dissolution of double Holliday junctions (Gari et al., 2008b; Karow et al., 2000; Willis et al., 2014; Wu and Hickson, 2003; Xue et al., 2015). Thus, BLM and, possibly, FANCM might act on post-replicative HJs to reduce the likelihood of their conversion to D-loops at stalled forks. Indeed, the Bloom’s homolog *SGS1* plays a role in postreplication repair at a Tus/*Ter* RFB in *S. cerevisiae* (Larsen et al., 2017).

Whatever its mechanism of formation, the established D-loop would also be a target for direct dissociation by FANCM/BLM (Bachrati et al., 2006; Gari et al., 2008a; Sun et al., 2008; Xue et al., 2015). In the context of FANCM/BLM dysfunction, failure to dissociate D-loops in a timely fashion might permit resumption of DNA synthesis and the establishment of BIR (**Figure 7C** and **7D**). This model is supported by *in vivo* studies in yeast: Sc*MPH1* (the *FANCM* homolog) promotes template switching in preference to processive BIR; in addition, *ScMPHl* and *SGSl* (the *BLM* homolog) confer a delay in the onset of BIR (Jain et al., 2016; Stafa et al., 2014).

The dramatic synergistic induction of TDs in cells lacking both FANCM and BRCA1 suggests that the two proteins act at different steps to suppress TD formation. If FANCM acts at the stalled fork to disallow BIR-mediated fork restart, BRCA1 may act upon a later step to suppress TD formation. We have suggested that BRCA1 suppresses TDs by promoting collapse of the nascent TD back to single copy status, through BRCA1’s role as a mediator of single strand annealing (Scully et al., 2019).

### Phenotype of ATPase-defective FANCM: synthetic lethality with *Brca1* mutation

We find that the ATPase (motor) function of FANCM is essential for all aspects of FANCM-mediated repair. A significant clue regarding the underlying mechanisms came from our ChIP analysis of FANCM at the Tus/*Ter* RFB. Strikingly, the FANCM ΔDEAH protein accumulates at Tus/*Ter* to levels ~2-fold higher than the wild type protein. Since *Fancm*^ΔDEAH^ was expressed at the same level as wild type *Fancm*, this finding suggests that the residence time of ATPase-defective FANCM at the stalled fork is prolonged compared to that of the wild type protein. ATP hydrolysis by FANCM may therefore contribute to the timely displacement of FANCM from the stalled fork. Conversely, a defect in ATP hydrolysis might “trap” FANCM at the stalled fork, with potential for neomorphic or antimorphic functions. Interestingly, small molecule inhibitors of poly(ADP ribose) polymerase (PARP), which are used in the treatment of *BRCA*-linked cancers, are thought to act, in part, by trapping PARP1 at sites of DNA damage (Murai et al., 2012).

Unexpectedly, we observed synthetic lethality between certain *Fancm* mutations and *Brca1* mutation. *Fancm*^Δ85/Δ^ (presumptively *Fancm* null) and *Fancm*^ΔDEAH^ cells were synthetic lethal with *Brca1* exon 11 deletion, while *Fancm*^ΔMM1^ and *Fancm*^ΔMM2^ cells were viable on this *Brca1* mutant background. It will be important to identify the mechanisms underlying synthetic lethality. This synthetic lethal relationship might explain why *FANCM* mutations are not observed in Group 1 TD *BRCA1*-linked breast or ovarian cancers—a puzzle, given the strong induction of cancer-promoting TDs that would occur in cells lacking both BRCA1 and FANCM (Menghi et al., 2018; Willis et al., 2017). If loss of *FANCM* ATPase function were lethal in *BRCA1* mutant cancers—as it is in *Brca1* mutant ES cells—this interaction could be exploited for cancer therapy, since the ATPase function of FANCM should, in principle, be “druggable”. Small molecule FANCM ATPase inhibitors might selectively kill *BRCA1* mutant cancer cells, while leaving surrounding *BRCA1*^+/−^ cells intact.

## Acknowledgements

We thank Drs. Andrew Deans, Hilda Pickett, Angelos Constantinou and Johannes Walter for helpful discussions. This work was supported by AACR fellowship 19-40-12-PAND (to A.P.), CDMRP grant BC160172 (to E.T.L. and R.S.) and NIH grants R01CA095175 and R01CA217991 (to R.S.).

## Author Contributions

A.P., N.A.W., R.E., F.M. and E.E.D. conducted the experiments; A.P., N.A.W., E.T.L. and R.S. designed the experiments. A.P., N.A.W. and R.S, wrote the paper.

## STAR Online Methods

### Molecular biology, siRNA and sgRNA oligos

C-terminal HA tagged Tus-F140 expression plasmid (pcDNA3β-MYC-NLS-Tus-F140A-3×HA) was derived from the parental vector (PMID29168504, PMID 24776801) and *6xTer* HR reporters were engineered by conventional cloning methods as previously described (PMID29168504, PMID 24776801). *Ter*-containing plasmids were amplified in JJC33 (Tus^−^) strains of *E. coli*. All plasmids used for transfection were prepared by endotoxin-free maxiprep (QIAGEN Sciences). All primers used for genotyping, RT-qPCR, sgRNA synthesis, and chromatin immunoprecipitation were purchased from Life Technologies. siRNA SMARTpools were purchased from Horizon Scientific/Dharmacon.

### Cell culture and the generation of mouse *Fancm* mutant cell lines

*Fancm* mutant cell lines were derived from a conditional *Brca1*^fl/exon11^ founder cell line carrying one *Brca1* allele in which much of exon 11 is replaced by the hygromycin resistance cassette, and one functionally wild type *Brca1* allele in which exon 11 is flanked by loxP elements and may be excised *Cre* recombinase. The founder cell line contains a single copy of the 6X *Ter*-HR reporter cassette, targeted to the *rosa26* locus and verified by southern blot (PMID 29168504, PMID 24776801, PMID 23994874). Mouse ES cells are routinely thawed onto plates coated with mouse embryonic fibroblast (MEF) feeders, maintained in ES medium on gelatinized plates, and regularly tested for mycoplasma infection by Myco-Alert assay (Lonza). *Fancm* mutant cell lines were generated using CRISPR/Cas9 mediated mutation of the *Fancm* locus. sgRNAs targeting Cas9 to the *Fancm* locus were transcribed *in vitro* using the Engen sgRNA Synthesis kit (New England Biolabs E3322S), purified using the RNA Clean and Concentrator Kit (Zymo Research, R1017), and verified by denaturing 10% TBE-urea PAGE. Cas9-sgRNA RNP was pre-assembled *in vitro* in OptiMEM by mixing Spy NLS Cas9 (New England Biolabs, M0646T) and purified sgRNA. Cells were co-transfected with either 0.45 μg Cas9(1.1) expression plasmid and 0.05 μg of each purified sgRNA using Lipofectamine 2000 (Invitrogen). After 72 hr, transfected cells were plated onto 6-cm dishes containing feeder MEFs without selection. Individual clones were picked for expansion between 9 and 14 days later and *Fancm* mutant clones were identified by PCR and confirmed by sequencing. *Fancm* targeting sgRNA sequences including PAM: *Fancm^Δ85/Δ85^* 5’ exon 2 (5’-GGT TCT TTT CCT GAC ACC GCA GG-3’); *Fancm^Δ85/Δ85^* 3’ exon 2 (5’-GAT GAA GCT CAT AAG GCA CTT GG-3’). *Fancm* exon 2 specific primers: *Fancm*^Δ85/Δ85^sense (5’-CTA CCT CAA GCT CCA GAG TCC TGG-3’); *Fancm*^Δ85/Δ85^, antisense (5’-AGT TCC CAT CAC TGA GAC TTA TTC C-3’).

### Generation of mouse *Fancm* hemizygous and in-frame mutant cell lines

*Fancm* domain and ATPase mutants were derived from a hemizygous (*Fancm*^+/−^) clone (**Supplemental Figure S2**). To generate *Fancm* clones in Brca1^fl/exon11^ 6X *Ter*-HR reporter cells, *Fancm* exons 2-23 were deleted by co-transfection with 30 nM preassembled Cas9-sgRNA RNP. RNP was preassembled *in vitro* in OptiMEM mixing Spy NLS Cas9 (New England Biolabs, M0646T) and purified sgRNA. *Fancn*^+/−^ clones which retained one wild type allele were identified by PCR and confirmed by PCR product sequencing. *Fancm* targeting sgRNA sequences including PAM: *Fancm* exon 2 (5’-GGT TCT TTT CCT GAC ACC GCA GG-3’); *Fancm* exon 23 (5’-GTT ACC AAA TGA TCT TAA CCA GG-3’). *Fancm* primers used to identify loss of exons 2-23: sense (5’-CTA CCT CAA GCT CCA GAG TCC TGG-3’); antisense (5’-AGG GAT GAC CTG AGG TTG TC-3’). *Fancm* primers specific for exons 2 and 23 break sites: exon 2 sense (5’-CTA CCT CAA GCT CCA GAG TCC TGG-3’); exon 2 antisense (5’-AGT TCC CAT CAC TGA GAC TTA TTC C-3’); exon 23 sense (5’-CAT GAG CCA GGA TAA AAA TAG TAA TTT C-3’); exon 23 antisense (5’-AGG GAT GAC CTG AGG TTG TC-3’).

To generate MM1 *(Fancm*^ΔMM1/−^), MM2 *(Fancm*^ΔMM2/−^), DEAH *(Fancm*^ΔDEAH/−^), MM3 *(Fancm*^ΔMM3/−^) mutants, the *Fancm*^+/−^ mutant was subjected to a second round of Cas9-sgRNA RNP-mediated targeted gene modification. Clonal candidates harboring the expected in-frame domain deletion were identified by PCR and confirmed by sequencing. *Fancm* targeting sgRNA sequences including PAM: MM1 5’-breaksite (ATG CTG ACA CTG TTA AAC AAA GG); MM1 3’-breaksite (GTG AAC AGC TCT TCT TCC AAT GG); MM2 5’-breaksite (CAA GAA GAG CTG AGG ACT GAC GG); MM2 3’-breaksite (TCT GAT AGG ACT CGC ACC CTG GG); DEAH 5’-breaksite (TGG TAA ATG ACC TTA CTA GAG GG); DEAH 3’-breaksite (ATG AAG CTC ATAA GGC ACT TGG G); MM3 5’-breaksite (AGA GGT TTC TGG AGA GGA TGT GG); MM3 3’-breaksite (TAA AGA TTC CTT TAC AGA TGA GG). *Fancm* primers specific for domain loss: MM1 sense (CTT GTT TGG TAG GGT GAA TGC A); MM1 antisense (GGG AGA ACG GGA TAA AAA TCT CT); MM2 sense (TAG ATG ATG ATT CTG AAC CTG AAG AC); MM2 antisense (TGT GCT CCT GAC TCT CTG CT); DEAH sense (CTA CCT CAA GCT CCA GAG TCC TGG); DEAH antisense (AGT TCC CAT CAC TGA GAC TTA TTC C); MM3 sense (CTG TCT TCA TGT CTC TTG TTT CT); MM3 antisense (AAT TCT AGA CAA TTT CTT TCT TGG C).

### Recombination assays

To measure HR repair frequencies, 1.6 x 10^5^ cells were co-transfected in suspension with 0.35 μg empty vector, pcDNA3β-myc NLS-I-SceI, or pcDNA3β-myc NLS-Tus and 20 pmol ON Targetplus siRNA SMARTpool using Lipofectamine 2000 (Invitrogen). GFP^+^RFP^−^, GFP^+^RFP^+^ and GFP^−^RFP^+^ frequencies were measured in duplicate 72 hr after transfection by using flow cytometry (Beckman Coulter CytoFlex LX). The total number of events that were scored for each sample was 3×10^5^−6×10^5^. Repair frequencies showed are corrected for background events and for transfection efficiency (55–85%). To measure transfection efficiency, parallel transfection was done with 0.05 μg wild-type GFP expression vector, 0.30 μ g control vector and 20 pmol siRNA. To deplete two gene targets, 10 pmol of each siRNA was used, while single depletion controls received 10 pmol of the target siRNA and 10 pmol of control luciferase siRNA. Data presented represent mean and error bars represent the s.e.m. of between 3 (n = 3) and 8 (n = 8) independent experiments (n values given in figure legends).

### Competition assays

To assess the mitomycin C (MMC) sensitivity of *Fancm* mutant clones, competition experiments were performed. 1.6 x 10^5^ cells were co-transfected in suspension with 0.45 μg empty vector and either 50 ng empty vector for each *Fancm* mutant clone or 50 ng GFP-expression plasmid for the parental wild type clone, using Lipofectamine 2000 (Invitrogen). 18 hours after transfection, cells were counted, and each *Fancm* mutant clone mixed and plated 1:5 with the GFP-marked parental wild type clone. 6 hours after cell plating, growth medium was replaced with media containing titrated doses of mitomycin C. Two days later, GFP+ frequencies were scored on a Beckman Coulter CytoFlex LX. Fold enrichment of cultures transiently co-transfected with GFP-expression plasmid normalized to 0 μg/mL mitomycin C control. Plots represent the mean of triplicate samples from three independent experiments (n = 3).

### Isolation of Chromatin-bound Proteins for FANCM western blot

Freshly harvested cell pellets were washed with 1x PBS and lysed in 5 volumes of chilled A1 buffer (50mM HEPES,140mM Nacl,1mMEDTA,10%glycerol,0.5% NP-40,0.25% TritonX-100,1mM DTT, and 1X protease inhibitor). Each lysate was centrifuged at 1,100 x g at 4 °C for 2 min and the supernatant was discarded. Pellets were resuspended by gentle pipetting in A1 buffer and samples were incubated for 10 minutes on ice. Each suspension was centrifuged at 1,100 x g at 4 °C for 2 min and the supernatant again discarded. Pellets were resuspended by gentle pipetting in 2 volumes of ice cold E2 buffer (10mM Tris-HCl pH-8,200mMNacl, 1mM EDTA and 0.5mM EGTA, and 1X Protease inhibitor). Each suspension was centrifuged at 1,100 x g at 4 °C for 2 min and the supernatant discarded. Pellets were resuspended by gentle pipetting with A1 buffer and incubated for 10 minutes on ice. Each suspension was centrifuged at 1,100 x g at 4 °C for 2 min and the supernatant discarded. Pellets were resuspended by gentle pipetting in 2 volumes of ice cold E3 buffer (500mM Tris-HCl pH-6.8, 500mMNacl and 1x Protease inhibitor. Each sample was sonicated in a water bath using Diagenode Bioruptor 300 with attached 4°C chiller cycling 30 sec on and 30 sec off on high setting for 5 min. Sonicated samples were centrifuged at 16,000xg for 15 min at 4°C. The supernatant corresponding to the chromatin fraction was transferred and subjected for western blot analysis using following antibodies; FANCM (Abcam ab95014,1:1000), Histone H3 (Abcam ab1791,1:1000).

### Chromatin immunoprecipitation assays

Chromatin immunoprecipitation was performed as described previously (Willis et al., 2018). Transfection of 2.0 x 10^5^ cells containing a single copy of a minimal cassette lacking redundant sequence harboring a single GFP allele interrupted by *6xTer-I-SceI* elements targeted to the *Rosa26* locus were performed with 0.5 μg empty vector, pcDNA3β-MYC NLS-I-SceI or pcDNA3β-MYC NLS-Tus-F140A-3xHA using Lipofectamine 2000 (Invitrogen). 10 million cells were collected 24 hours after transfection and fixed in 1% formaldehyde supplemented serum free media. Fixation was quenched by addition of glycine to 125 mM. Cells were lysed in lysis buffer (0.1% SDS, 20 mM EDTA, 50 mM Tris pH 8.1) containing protease inhibitor (Roche protease inhibitor, Roche 13539320). Chromatin shearing to ~500 bp was accomplished using Diagenode Bioruptor 300 with attached 4°C chiller for 20 cycles, 15 seconds on and 30 seconds off at medium setting. To avoid non-specific binding to protein A/G beads, 100 μl lysates for each ChIP sample were precleared by the addition of 10 μl activated Magna ChIP magnetic beads (Millipore Sigma, 16-663) in ChIP dilution buffer (1% Triton-X-100, 2mM EDTA, 150mM NaCl, 20mM Tris pH 8.1) and incubation for 1 hr at 4°C with gentle mixing. After removal of beads by magnet, for each immunoprecipitation, 3 μl of anti-FANCM (Abcam ab95014), 3 μl of anti-FANCA (Abcam ab97578), 3 μl of anti FANCL (Abcam ab94458), 3 μl of Rad51(Abcam, ab176458), or 5 μl of anti-Blm (Bethyl A 300-110A) was added and mixed for 12 hours at 4°C followed by addition of 10 μl activated Magna ChIP beads and mixing for 16 hours at 4°C. Beads were washed six times in ice-cold ChIP RIPA buffer (50mM HEPES pH 7.6, 1mM EDTA, 7 mg/mL sodium deoxycholate, 1% NP-40) followed by two washes in ice-cold TE (<u>10 mM Tris pH8.0, 1 mM EDTA</u>). Crosslinks were reversed and DNA was eluted by incubation in 100 μL Elution buffer (1% SDS, 200mM sodium bicarbonate, 5.6 μg/mL RNAse A) overnight at 65°C. Protein was removed by proteinase K digest for 30 min at 55°C. Released DNA was purified by Qiagen PCR Purification column (Qiagen, 28106) and analyzed by qPCR on an ABI Prism 7300 or QuantStudio3 using SYBR Green (Applied Biosystems, 4368702). Primers for qPCR: +109 bp sense (TCC GGA TAG GGA TAA CAG GGT A); +109 bp antisense (GTC GGC CAT GAT ATA GAC GTT G); +309 bp sense (AGC TCG CCG ACC ACT AC); +309 bp antisense (TCC AGC AGG ACC ATG TGA T); +921 bp sense (GGA CAA GAC TTCCCA CAG ATT); +921 bp antisense (GAG GCG GAT CAC AAG CAA TAA T); +1.6 kb sense (TCC ACA TTT GGG CCT ATT CTC); +1.6 kb antisense (CAA TAA TGA AAT ATA CCT TTT AAT GTC T); 128 bp sense (GAG CGC ACC ATC TTC TTC A); 128 bp antisense (TCC CTA CGA TGC CCT TCA); 350 bp sense (CTG GAC GGC GAC GTA AAC)); 350 bp antisense (CGG TGG TGC AGA TGA ACT T); 900 bp-sense (TCT GGA GCA TGC GCT TTA G); 900 bp antisense (CTA AAG CGC ATG CTC CA GA); 1.4kb sense (CCA CTG CCC TTG TGA CTA AA); 1.4kb antisense (AGG CTA CAC CAA CGT CAA TC). Data are presented as the mean calculated from three independent experiments (n = 3) normalized against untreated controls (empty vector) and control locus (beta-actin) using the 2^−ΔΔCT^ method (REF).

### RT-qPCR analysis

RNA extraction was performed using QIAGEN RNeasy Mini Kit (QIAGEN Sciences, Maryland, MD). All analyses of *Gapdh* and genes of interest were performed using an Applied Biosystems 7300 Real time PCR System using Power SYBR Green RNA-to CT™ 1-Step Kit (Applied Biosystems, Foster City, CA).Primers for RT-PCR: *Gapdh* sense (CGT CCC GTA GAC AAA ATG GT); *Gapdh* antisense (TCG TTG ATG GCA ACA ATC TC); *Fancm* sense (GTC GTT ATC CTC GCT GAA GG); *Fancm* antisense (TTT GTT GGA CTG ACT CTG ATT ATA TGT); MM1 sense (CTG TTA AACAAA GGG ATT CTA AAT); MM1 antisense (GAT ACA GAT TTC TCA TCA CTG A); MM2 sense (TCG TTG TAG TTC GGG TTC AGA); MM2 antisense (AGT GTT CAA CTT CAG TGC GCC); MM3 sense (TCC TGA GCA AGA TGA AAA CTA T); MM3 antisense (TTA TAT CGT ATG TCC TCA TCT GTA A); DEAH sense (TGG CTG AAA TGA CAG GTT CAA CT); DEAH antisense (GCC TTA TGA GCT TCA TCC ACC). mRNA was measured in technical triplicates. Target gene expression level was normalized to *Gapdh* using the 2^−ΔCT^ method.

### Statistical methods

Data shown represents the arithmetic mean and error bars represent the standard error of the mean (s.e.m.) of between three (n = 3) and nine (n = 9) independent experiments (n values given in figure legends). Each Figure Legend reports the number of independent experiments (n) that were performed to generate the data presented. Data points for each independent experiment were typically collected as the mean of technical duplicates. This mean value was taken as the solitary data point for that individual experiment. The arithmetic mean of samples collected for groups of independent experiments for repair frequency statistical analysis, was calculated and data points for each independent experiment used to calculate the mean and s.e.m., calculated as standard deviation/√ n, (n indicates the number of independent experiments). Differences between sample pairs repair frequencies and fold enrichment for ChIP were analyzed by Student’s two-tailed unpaired t-test, assuming unequal variance. One-way ANOVA statistical analysis of greater than three samples was performed when indicated P-values are indicated in the figure and or figure legends. No statistical methods were used to predetermine sample size. The experiments were not randomized, and investigators were not blinded to allocation during experiments and outcome assessment. All statistics were performed using GraphPad Prism v7.0d software.

## Supplemental Figure Legends

**Figure S1.**
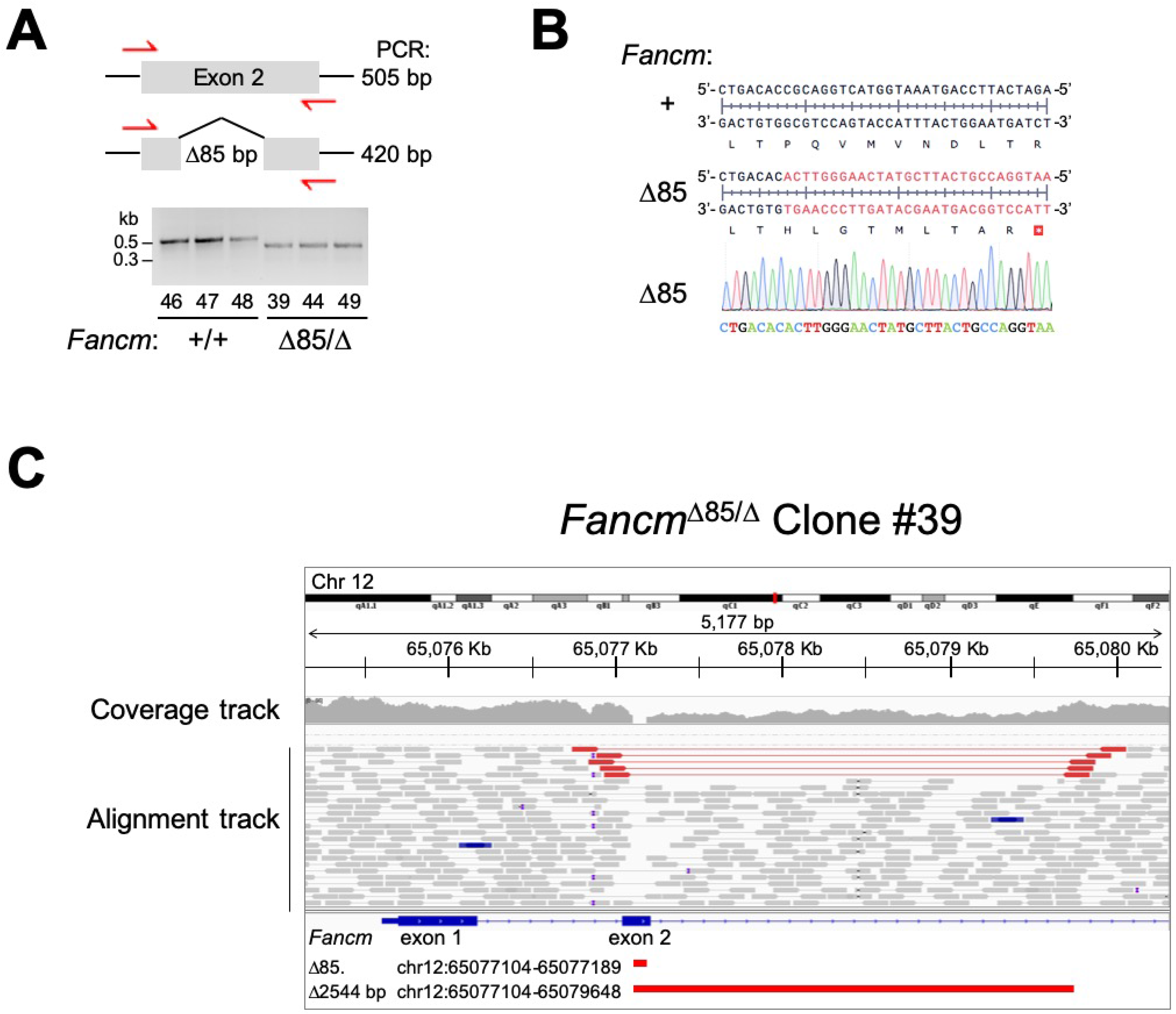
Genotype of *Fancm*^Δ85/Δ^ *Ter*-HR reporter clone #39. Related to Figure 1. **A.** PCR primers for detection of 85 bp frame-shift *Fancm*^Δ85^ allele. Red half-arrow heads: genotyping primers. Gel: PCR products of *Fancm*^+/−^ and *Fancm*^Δ85/Δ^ clones. **B.** Primary DNA sequencing chromatogram of PCR product from a *Fancm*^Δ85/Δ^ clone harboring the *Fancm*^Δ85^ allele. **C.** Whole genome sequencing reads spanning *Fancm* exons 1 and 2 in *Fancm*^Δ85/Δ^ clone #39. Note how 85 bp deletion within exon 2 results in zero coverage. Red gapped reads in alignment track identify 2544 bp deletion in second (*Fancm*^Δ^) allele, overlapping the 85 bp deletion of the *Fancm*^Δ85^ allele. *Fancm* exons are shown as blue bars below alignment track. Red bars below exons identify exact chromosome positions of the two deletions.

**Figure S2.**
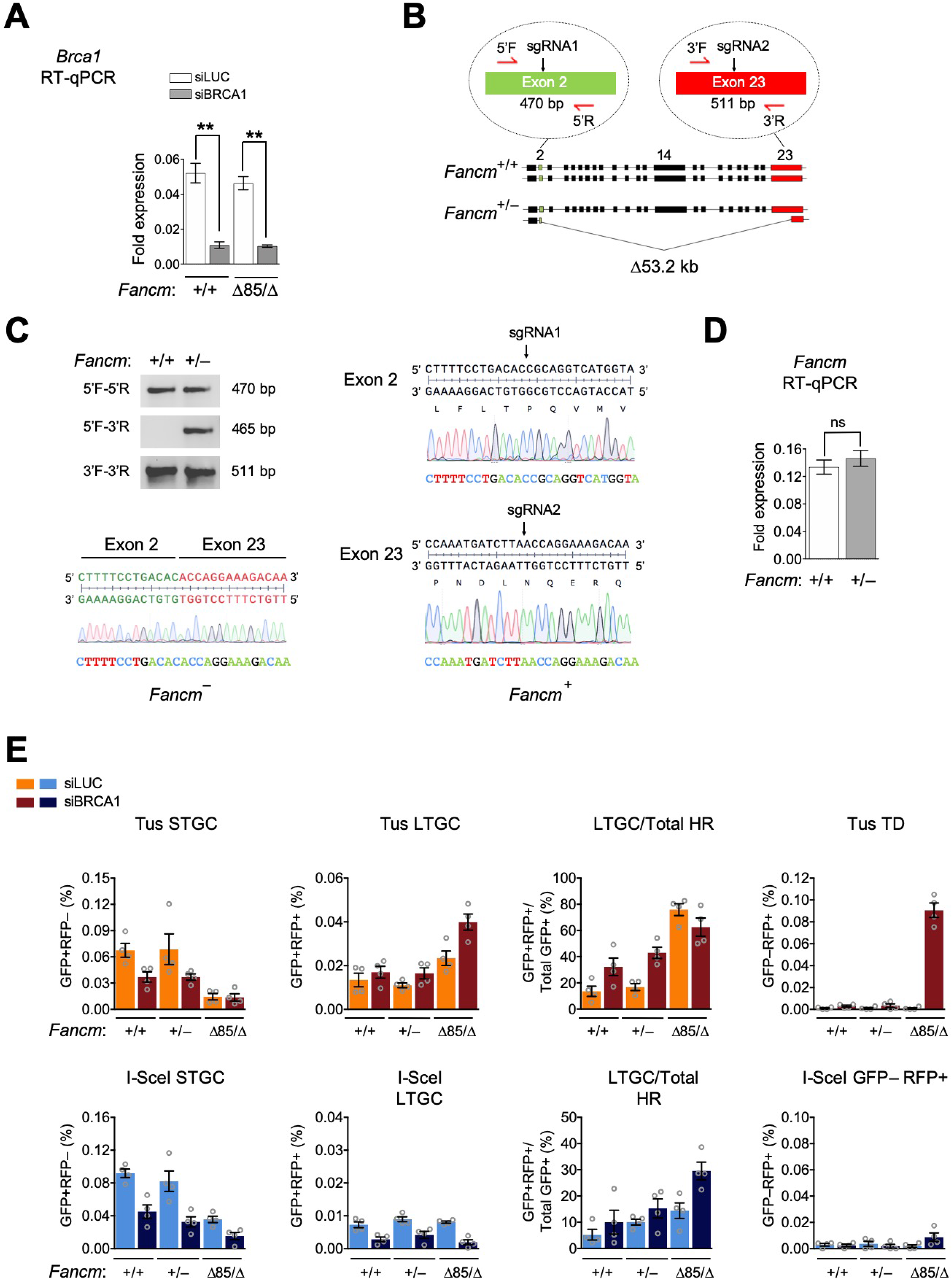
*Fancm* hemizygous cells retain wild type stalled fork repair phenotypes. Related to Figure 2. **A.** RT qPCR analysis of *Brca1* mRNA in *Fancm*^+/+^ or *Fancm*^Δ85/Δ^ clones treated with siRNAs shown. Data shows mean of three independent experiments (i.e., n=3) normalized to *Gapdh* mRNA using the 2^−ΔΔCT^ method and analyzed by Student’s *t*-test. Error bars: standard deviation. **: P<0.01. **B.** Gene modification strategy to generate hemizygous *Fancm*^+/−^ clones using *S.p*. Cas9 with dual sgRNAs targeting exons 2 and 23. The *Fancm*^−^ allele harbors the expected loss of 53.2 kb intervening sequence in the *Fancm* gene. Red half arrowheads: PCR and sequencing primers specific to Exons 2 and 23. Predicted PCR product size for *Fancm*^+^ wild type allele shown. **C.** Gel: PCR products of *Fancm*^+/+^ and *Fancm* clones. Below: Primary DNA sequencing chromatogram of *Fancm*^−^ allele PCR product verifies exon2 (green) to exon 23 (red) breakpoint. Right: Primary DNA sequencing chromatograms of *Fancm*^+^ allelic PCR products from the same clone verify wild type sequence of *Fancm*^+^ allele at sgRNA target sites. **D.** RT qPCR analysis of *Fancm* mRNA in *Fancm*^+/+^ or *Fancm*^+/−^ clones. Data normalized to *Gapdh* using the 2^−ΔCT^ method (n=3) using Student’s *t*-test. Error bars: standard deviation. ns: not significant. **E.** Tus/*Ter*- and I-SceI-induced repair in *Fancm*^+/+^, *Fancm*^+/+^, and *Fancm*^Δ85/Δ^ *Ter*-HR reporter clones co-transfected with Tus or I-SceI expression plasmdis and siRNAs as shown. Data shows mean values (n=4). Error bars: s.e.m. *Fancm*^Δ85/Δ^ cells serve as control for *Fancm* mutant phenotype. All repair outcomes for *Fancm*^+/−^ *vs. Fancm* are not significant by Student’s *t*-test.

**Figure S3.**
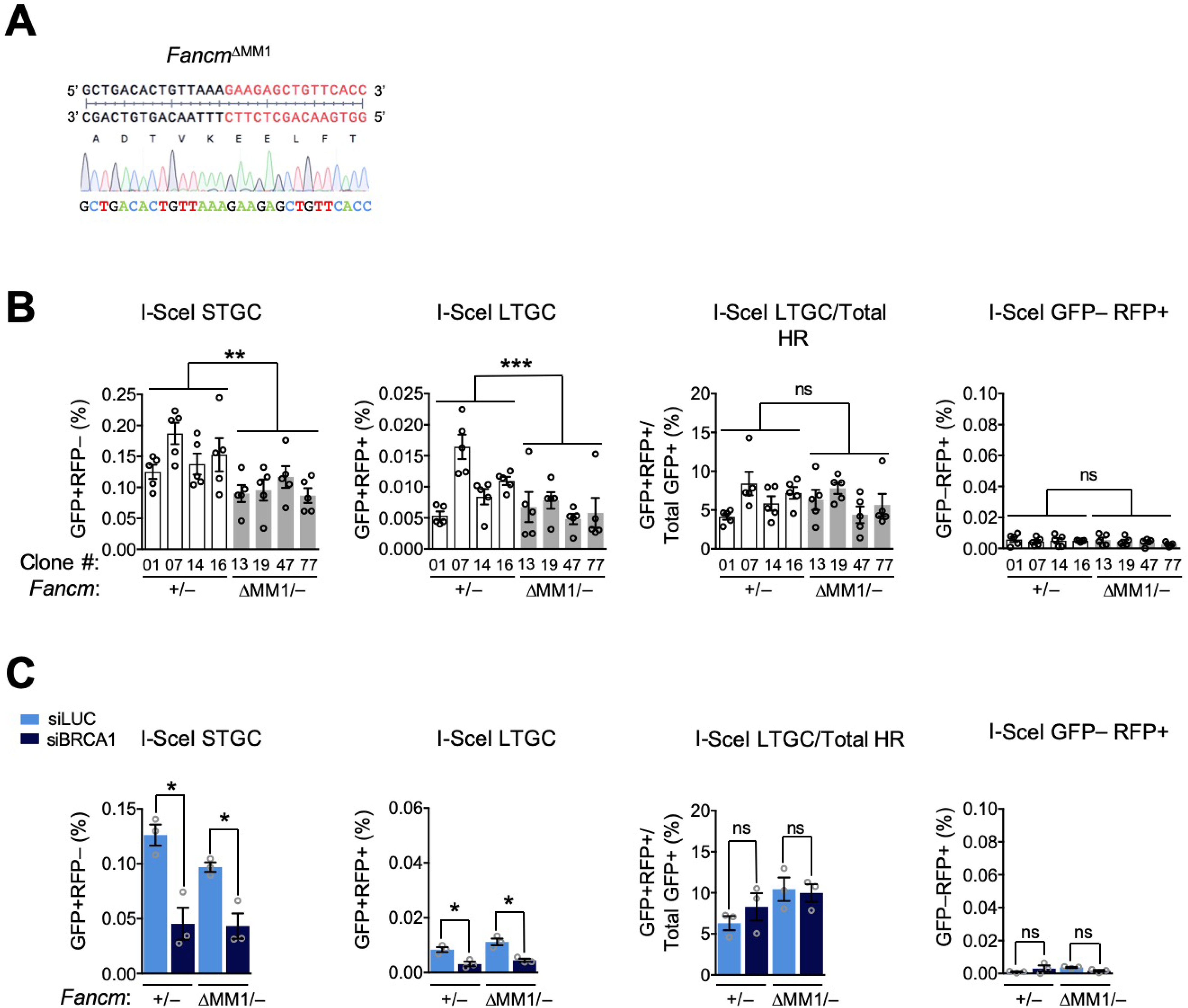
The FANCM-FA core complex interaction mediates I-SceI-induced HR. Related to Figure 3. A. Primary DNA sequencing chromatogram from a representative *Fancm*^ΔMM1/−^ clone indicates in-frame 366 bp deletion of MM1 coding sequence within the residual *Fancm* allele. **B.** I-SceI-induced HR in four *Fancm*^+/−^ (white bars) clones *vs*. four *Fancm*^ΔMM1/−^ (gray bars) clones. Data obtained from same experiments as in **Figure 3F**. Data shows mean values (n=5). Error bars: s.e.m. **: P < 0.01, ***: P < 0.001 by one-way ANOVA. ns: not significant. **C.** I-SceI-induced HR in *Fancm*^+/−^ *vs. Fancm*^ΔMM1/−^ clones co-transfected with I-SceI expression plasmid and siRNAs as shown. Data obtained from same experiments as in **Figure 3G**. Data shows mean values, n=3. Error bars: s.e.m. *: P < 0.05, by Student’s *t*-test. ns: not significant.

**Figure S4.**
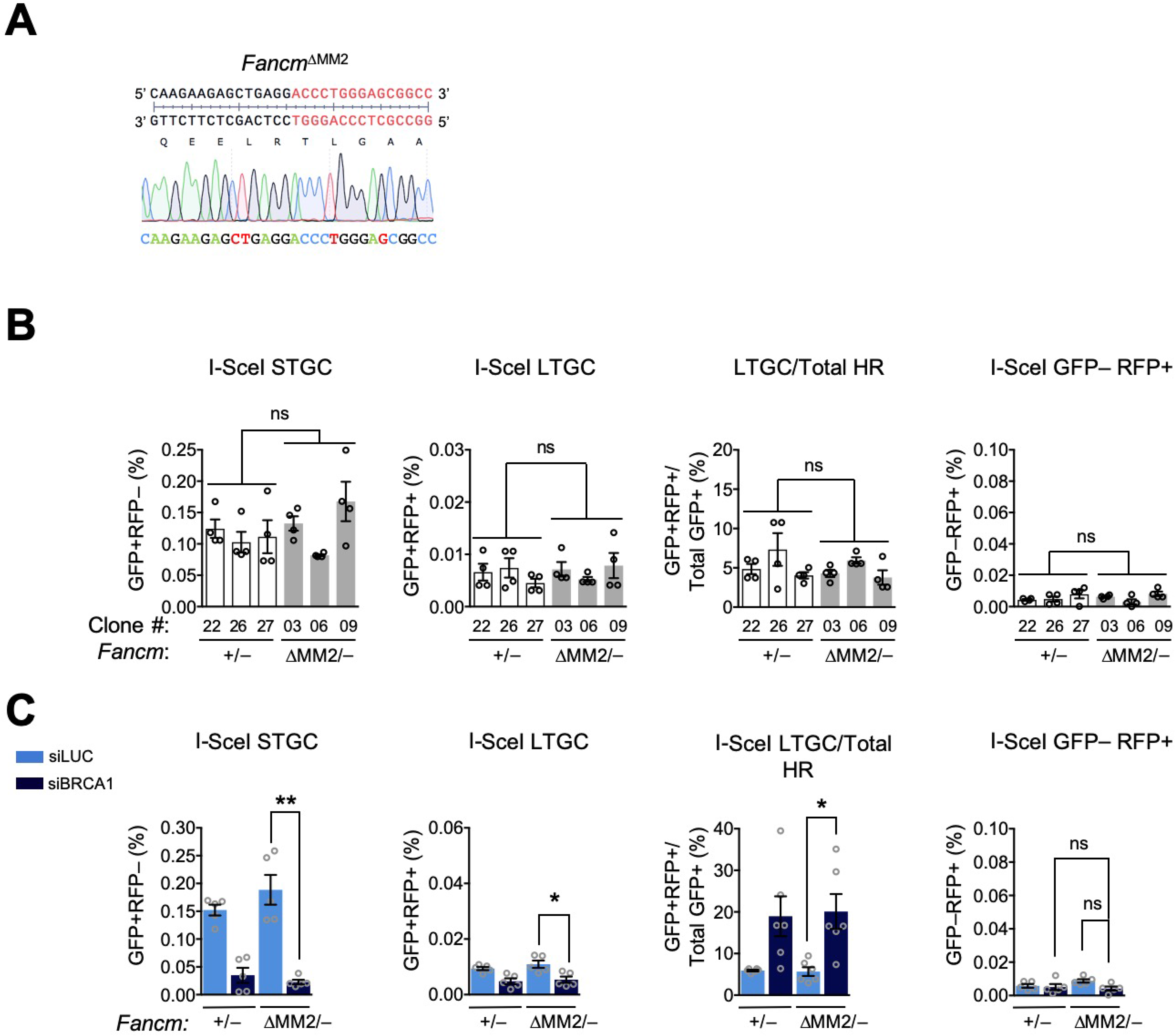
No impact of FANCM MM2 domain on DSB-induced HR. Related to Figure 4. **A.** Primary DNA sequencing chromatogram from a representative *Fancm*^ΔMM2/−^ clone indicates in-frame 114 bp deletion of MM2 coding sequence within the residual *Fancm* allele. **B.** I-SceI-induced HR in three *Fancm*^+/−^ (white bars) clones *vs*. three *Fancm*^ΔMM2/−^ (gray bars) clones. Data obtained from same experiments as in **Figure 4F**. Data shows mean values (n=4). Error bars: s.e.m. ns: not significant by one-way ANOVA. **C.** I-SceI-induced HR in *Fancm*^+/−^ *vs. Fancm*^ΔMM2/−^ clones co-transfected with I-SceI expression plasmid and siRNAs as shown. Data obtained from same experiments as in **Figure 4G**. Data shows mean values, n=5. Error bars: s.e.m. *: P < 0.05, **: P < 0.01 by Student’s *t*-test. ns: not significant.

**Figure S5.**
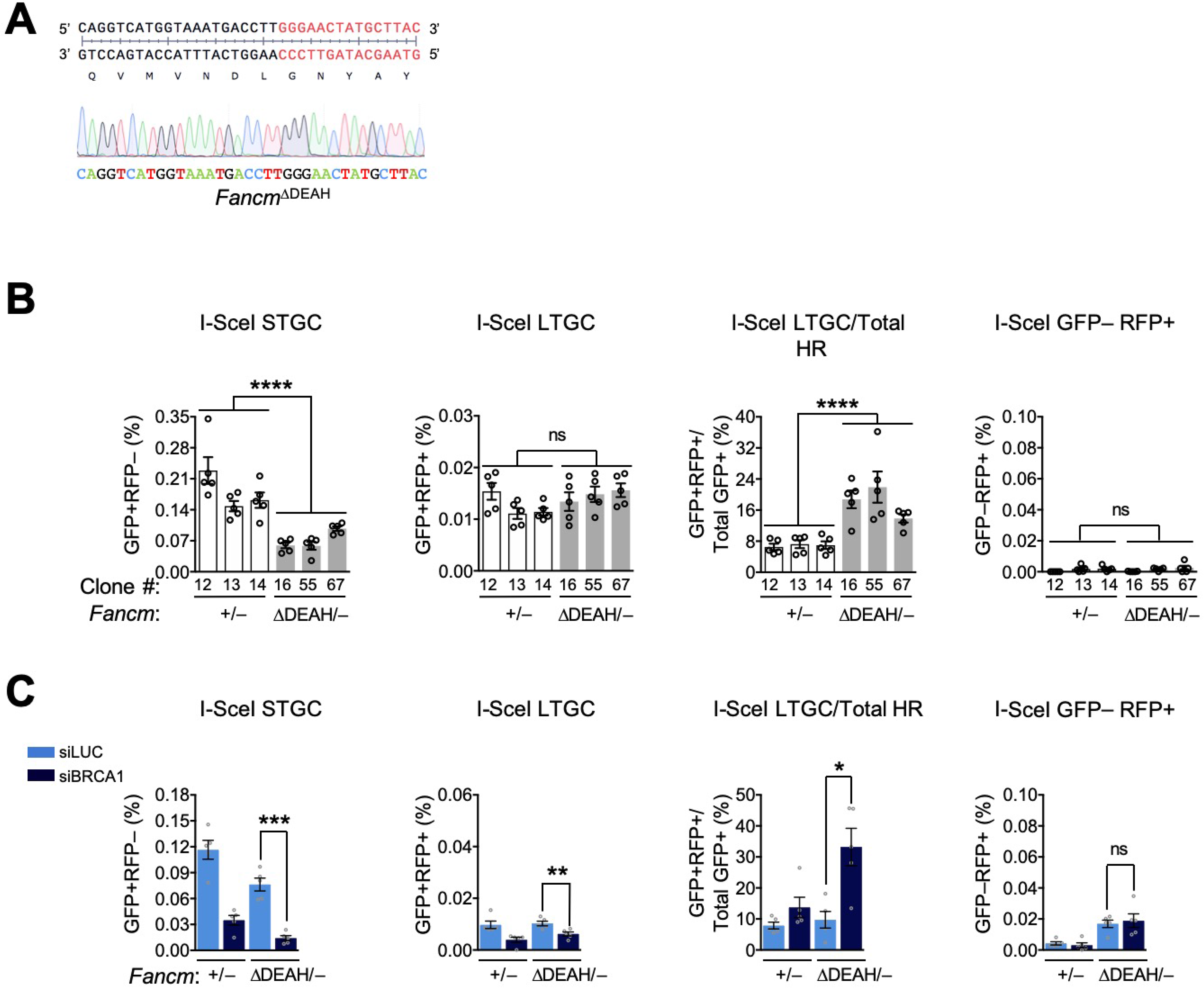
FANCM ATP hydrolysis mutant is defective for DSB-induced HR. Related to Figure 5. **A.** Primary DNA sequencing chromatogram from a representative *Fancm*^ΔDEAH/−^ clone indicates in-frame 66 bp deletion of sequence encoding the DEAH box within the residual *Fancm* allele. **B.** I-SceI-induced HR in three *Fancm* (white bars) clones *vs*. three *Fancm*^ΔDEAH/−^ (gray bars) clones. Data obtained from same experiments as in **Figure 5E**. Data shows mean values (n=5). Error bars: s.e.m. ****: P < 0.0001 by one-way ANOVA. ns: not significant. **C.** I-SceI-induced HR in *Fancm*^+/−^ *vs. Fancm*^ΔDEAH/−^ clones co-transfected with I-SceI expression plasmid and siRNAs as shown. Data obtained from same experiments as in **Figure 5F**. Data shows mean values, n=5. Error bars: s.e.m. *: P < 0.05, **: P < 0.01, ***: P < 0.001 by Student’s *t*-test. ns: not significant.

**Figure S6.**
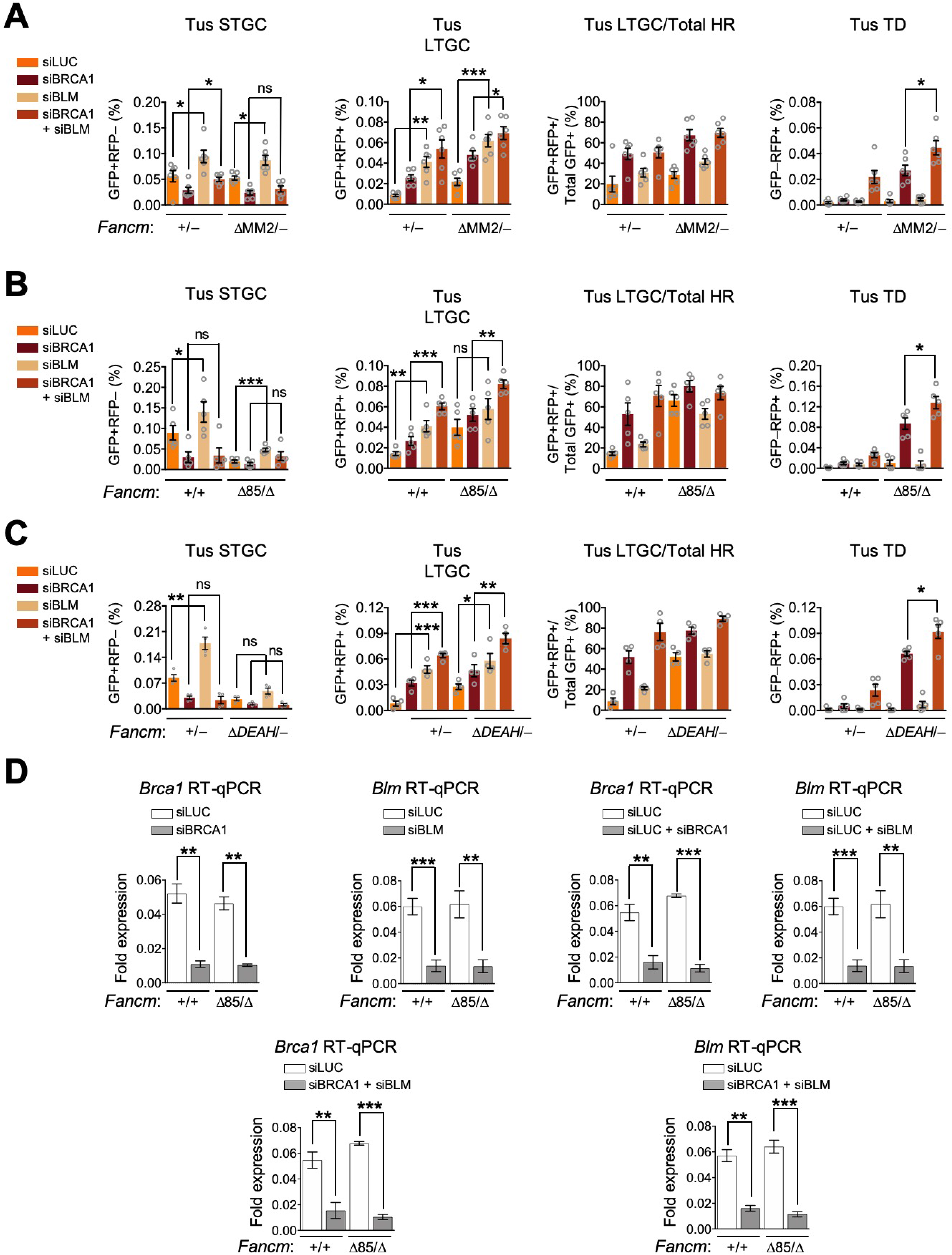
Stalled fork repair functions of BLM are *Fancm*-independent. See also Figure S6. **A.** Tus/*Ter*-induced repair in *Fancm*^+/−^ and *Fancm*^ΔMM22−^ clones co-transfected with Tus expression plasmid and siRNAs as indicated. Data shows mean values, n=6. Error bars: s.e.m. *: P<0.05, **: P < 0.01, ***: P < 0.001 by Student’s *t*-test. ns: not significant. **B.** Tus/*Ter*-induced repair in *Fancm^+/+^* and *Fancm*^Δ85/Δ^ clones co-transfected with Tus expression plasmid and siRNAs as indicated (n=5). Data analysis as in panel A. **C.** Tus/*Ter*-induced repair in *Fancm*^+/−^ and *Fancm*^ΔDEAH/−^ clones co-transfected with Tus expression plasmid and siRNAs as indicated (n=5). Data analysis as in panel A. **D.** RT qPCR analysis of *Brca1* or *Blm* mRNA in *Fancm*^+/−^, *Fancm*^Δ85/Δ^, or *Fancm*^ΔDEAH/−^ clones treated with siRNAs as shown. Mean fold difference calculated from three independent experiments (n=3). Data normalized to *Gapdh* using the 2^−ΔCT^ method (n=3) using Student’s *t*-test. **: P < 0.01, ***: P < 0.001. Error bars: standard deviation. ns: not significant.

**Figure S7.**
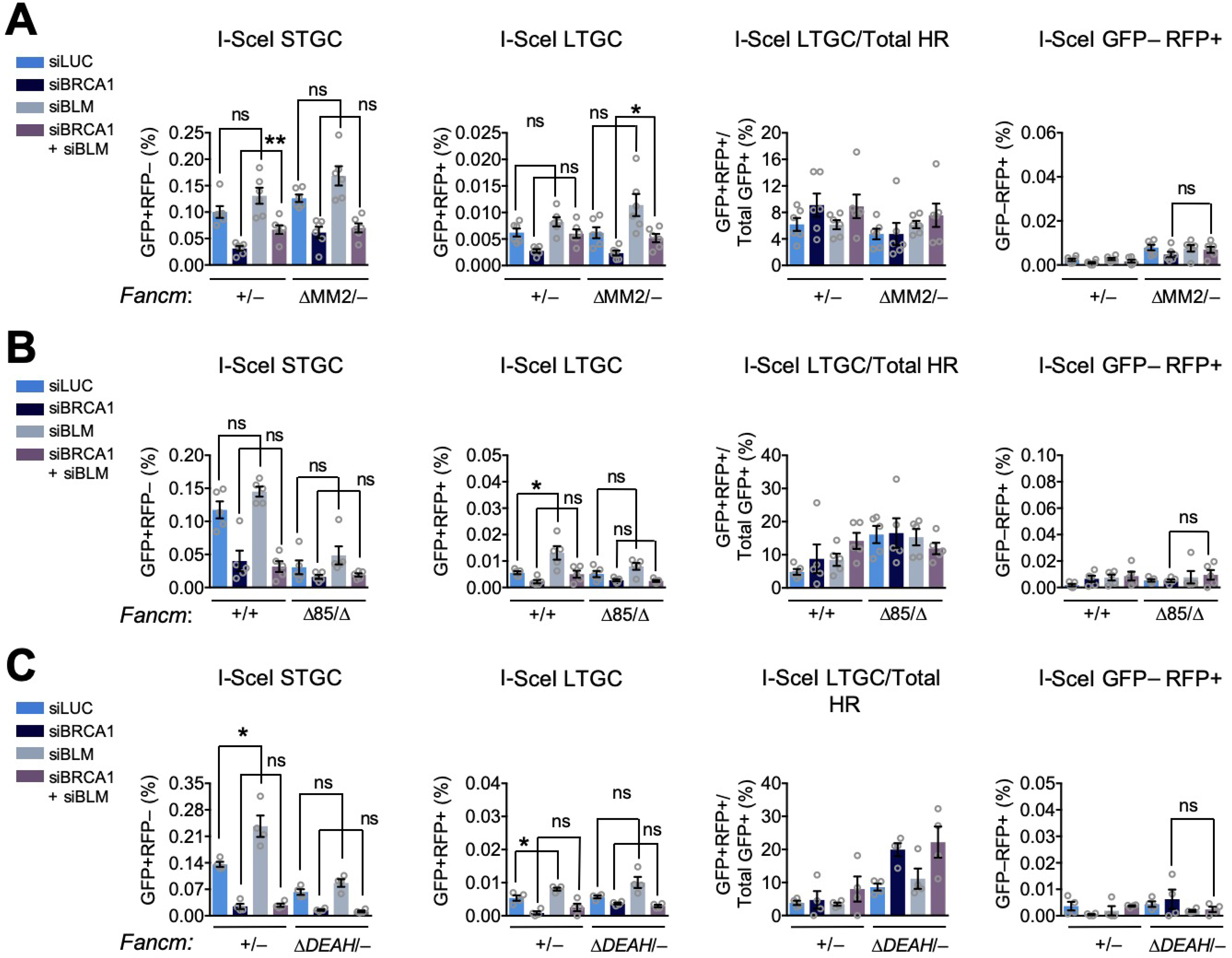
BLM depletion has minimal impact on DSB induced HR. Related to Figure S6. **A.** I-SceI-induced HR in *Fancm*^+/−^ and *Fancm*^ΔMM22/−^ clones co-transfected with I-SceI expression plasmid and siRNAs as indicated. Data shows mean values, n=6. Error bars: s.e.m. *: P<0.05, by Student’s *t*-test. ns: not significant. Data obtained from same experiments as in **Figure S6A**. **B.** I-SceI-induced repair in *Fancm*^+/+^ and *Fancm*^Δ85/Δ^ clones co-transfected with I-SceI expression plasmid and siRNAs as indicated (n=5). Data analysis as in panel A. Data obtained from same experiments as in **Figure S6B**. **C.** I-SceI-induced repair in *Fancm*^+/+^ and *Fancm*^ΔDEAH/−^ clones co-transfected with I-SceI expression plasmid and siRNAs as indicated (n=5). Data analysis as in panel A. Data obtained from same experiments as in **Figure S6C**.

**Figure S8.**
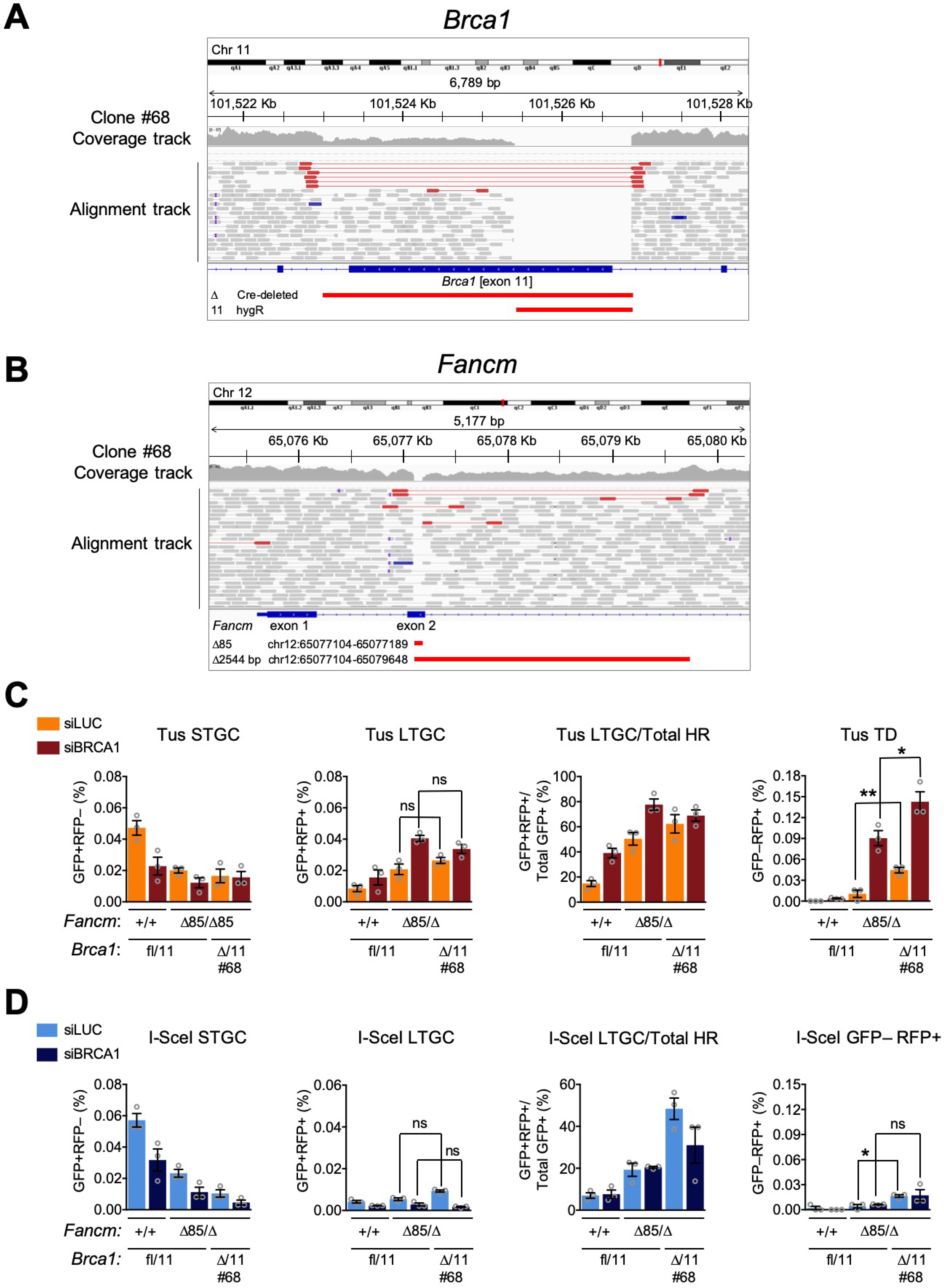
Analysis of solitary viable *Fancm*^Δ85/Δ^ *Brca1*^Δ/11^ clone #68. Related to Figure 6. **A.** Whole genome sequencing reads spanning *Brca1* exons 10, 11 and 12 in the solitary viable *Fancm*^Δ85/Δ^ *Brca1*^Δ/11^ clone #68. *Brca1* exons shown in blue bars beneath alignment track. *Fancm* exons are shown as blue bars below alignment track. Red bars below exons identify positions of the two deletions within exon 11. **B.** Whole genome sequencing reads spanning *Fancm* exons 1 and 2 in *Fancm*^Δ85/Δ^ clone #39. Elements as in Figure S1C. **C.** Tus/*Ter*-induced repair in *Fancm*^+/+^ or Cre-transduced *Fancm*^Δ85/Δ^ clones having retained or deleted (clone #68) *Brca1* exon 11, co-transfected with Tus-expression plasmid and siRNAs as shown. Note induction of TDs in *Fancm*^Δ85/Δ^ *Brca1*^Δ/11^ clone in absence of BRCA1 depletion. Data shows mean values, n=3. Error bars: s.e.m. *: P<0.05, **: P < 0.01 by Student’s *t*-test. ns: not significant. **D.** I-SceI-induced repair in the same experiments as in panel C.

## References

Adamo, A., Collis, S.J., Adelman, C.A., Silva, N., Horejsi, Z., Ward, J.D., Martinez-Perez, E., Boulton, S.J., and La Volpe, A. (2010). Preventing nonhomologous end joining suppresses DNA repair defects of Fanconi anemia. Mol Cell 39, 25–35.

Amunugama, R., Willcox, S., Wu, R.A., Abdullah, U.B., El-Sagheer, A.H., Brown, T., McHugh, P.J., Griffith, J.D., and Walter, J.C. (2018). Replication Fork Reversal during DNA Interstrand Crosslink Repair Requires CMG Unloading. Cell reports 23, 3419–3428.

Bachrati, C.Z., Borts, R.H., and Hickson, I.D. (2006). Mobile D-loops are a preferred substrate for the Bloom’s syndrome helicase. Nucleic Acids Res 34, 2269–2279.

Bakker, S.T., van de Vrugt, H.J., Rooimans, M.A., Oostra, A.B., Steltenpool, J., Delzenne-Goette, E., van der Wal, A., van der Valk, M., Joenje, H., te Riele, H., et al. (2009). Fancm-deficient mice reveal unique features of Fanconi anemia complementation group M. Human molecular genetics 18, 3484–3495.

Bogliolo, M., Bluteau, D., Lespinasse, J., Pujol, R., Vasquez, N., d’Enghien, C.D., Stoppa-Lyonnet, D., Leblanc, T., Soulier, J., and Surrallés, J. (2018). Biallelic truncating FANCM mutations cause early-onset cancer but not Fanconi anemia. Genetics in medicine: official journal of the American College of Medical Genetics 20, 458–463.

Castéra, L., Harter, V., Muller, E., Krieger, S., Goardon, N., Ricou, A., Rousselin, A., Paimparay, G., Legros, A., Bruet, O., et al. (2018). Landscape of pathogenic variations in a panel of 34 genes and cancer risk estimation from 5131 HBOC families. Genetics in medicine: official journal of the American College of Medical Genetics 20, 1677–1686.

Catucci, I., Osorio, A., Arver, B., Neidhardt, G., Bogliolo, M., Zanardi, F., Riboni, M., Minardi, S., Pujol, R., Azzollini, J., et al. (2018). Individuals with FANCM biallelic mutations do not develop Fanconi anemia, but show risk for breast cancer, chemotherapy toxicity and may display chromosome fragility. Genetics in medicine: official journal of the American College of Medical Genetics 20, 452–457.

Chandramouly, G., Kwok, A., Huang, B., Willis, N.A., Xie, A., and Scully, R. (2013). BRCA1 and CtIP suppress long-tract gene conversion between sister chromatids. Nat Commun 4, 2404.

Ciccia, A., and Elledge, S.J. (2010). The DNA damage response: making it safe to play with knives. Mol Cell 40, 179–204.

Ciccia, A., Ling, C., Coulthard, R., Yan, Z., Xue, Y., Meetei, A.R., Laghmani el, H., Joenje, H., McDonald, N., de Winter, J.P., et al. (2007). Identification of FAAP24, a Fanconi anemia core complex protein that interacts with FANCM. Mol Cell 25, 331–343.

Collis, S.J., Ciccia, A., Deans, A.J., Horejsi, Z., Martin, J.S., Maslen, S.L., Skehel, J.M., Elledge, S.J., West, S.C., and Boulton, S.J. (2008). FANCM and FAAP24 function in ATR-mediated checkpoint signaling independently of the Fanconi anemia core complex. Mol Cell 32, 313–324.

Cortez, D. (2019). Replication-Coupled DNA Repair. Mol Cell 74, 866–876.

Crismani, W., Girard, C., Froger, N., Pradillo, M., Santos, J.L., Chelysheva, L., Copenhaver, G.P., Horlow, C., and Mercier, R. (2012). FANCM limits meiotic crossovers. Science 336, 1588–1590.

Deans, A.J., and West, S.C. (2009). FANCM connects the genome instability disorders Bloom’s Syndrome and Fanconi Anemia. Mol Cell 36, 943–953.

Deans, A.J., and West, S.C. (2011). DNA interstrand crosslink repair and cancer. Nature reviews 11, 467–480.

Duxin, J.P., and Walter, J.C. (2015). What is the DNA repair defect underlying Fanconi anemia? Current opinion in cell biology 37, 49–60.

Figlioli, G., Kvist, A., Tham, E., Soukupova, J., Kleiblova, P., Muranen, T.A., Andrieu, N., Azzollini, J., Balmaña, J., Barroso, A., et al. (2020). The Spectrum of FANCM Protein Truncating Variants in European Breast Cancer Cases. Cancers 12.

Gari, K., Decaillet, C., Delannoy, M., Wu, L., and Constantinou, A. (2008a). Remodeling of DNA replication structures by the branch point translocase FANCM. Proc Natl Acad Sci U S A, 105, 16107–16112.

Gari, K., Decaillet, C., Stasiak, A.Z., Stasiak, A., and Constantinou, A. (2008b). The Fanconi anemia protein FANCM can promote branch migration of Holliday junctions and replication forks. Mol Cell 29, 141–148.

Hodskinson, M.R., Silhan, J., Crossan, G.P., Garaycoechea, J.I., Mukherjee, S., Johnson, C.M., Schärer, O.D., and Patel,K.J. (2014). Mouse SLX4 is a tumor suppressor that stimulates the activity of the nuclease XPF-ERCC1 in DNA crosslink repair. Mol Cell 54, 472–484.

Huang, J., Liu, S., Bellani, M.A., Thazhathveetil, A.K., Ling, C., de Winter, J.P., Wang, Y., Wang, W., and Seidman, M.M. (2013). The DNA translocase FANCM/MHF promotes replication traverse of DNA interstrand crosslinks. Mol Cell 52, 434–446.

Jain, S., Sugawara, N., Mehta, A., Ryu, T., and Haber, J.E. (2016). Sgs1 and Mph1 Helicases Enforce the Recombination Execution Checkpoint During DNA Double-Strand Break Repair in Saccharomyces cerevisiae. Genetics 203, 667–675.

Jalan, M., Oehler, J., Morrow, C.A., Osman, F., and Whitby, M.C. (2019). Factors affecting template switch recombination associated with restarted DNA replication. eLife 8.

Karow, J.K., Constantinou, A., Li, J.L., West, S.C., and Hickson, I.D. (2000). The Bloom’s syndrome gene product promotes branch migration of holliday junctions. Proc Natl Acad Sci U S A 97, 6504–6508.

Kim, H., and D’Andrea, A.D. (2012). Regulation of DNA cross-link repair by the Fanconi anemia/BRCA pathway. Genes Dev 26, 1393–1408.

Klein Douwel, D., Boonen, R.A., Long, D.T., Szypowska, A.A., Raschle, M., Walter, J.C., and Knipscheer, P. (2014). XPF-ERCC1 acts in Unhooking DNA interstrand crosslinks in cooperation with FANCD2 and FANCP/SLX4. Mol Cell 54, 460–471.

Knipscheer, P., Raschle, M., Smogorzewska, A., Enoiu, M., Ho, T.V., Scharer, O.D., Elledge, S.J., and Walter, J.C. (2009). The Fanconi anemia pathway promotes replication-dependent DNA interstrand cross-link repair. Science 326, 1698–1701.

Knoll, A., Higgins, J.D., Seeliger, K., Reha, S.J., Dangel, N.J., Bauknecht, M., Schropfer, S., Franklin, F.C., and Puchta, H. (2012). The Fanconi anemia ortholog FANCM ensures ordered homologous recombination in both somatic and meiotic cells in Arabidopsis. Plant Cell 24, 1448–1464.

Kosicki, M., Tomberg, K., and Bradley, A. (2018). Repair of double-strand breaks induced by CRISPR-Cas9 leads to large deletions and complex rearrangements. Nature biotechnology 36, 765–771.

Langevin, F., Crossan, G.P., Rosado, I.V., Arends, M.J., and Patel, K.J. (2011). Fancd2 counteracts the toxic effects of naturally produced aldehydes in mice. Nature 475, 53–58.

Larsen, N.B., Liberti, S.E., Vogel, I., Jorgensen, S.W., Hickson, I.D., and Mankouri, H.W. (2017). Stalled replication forks generate a distinct mutational signature in yeast. Proc Natl Acad Sci U S A 114, 9665–9670.

Long, D.T., Raschle, M., Joukov, V., and Walter, J.C. (2011). Mechanism of RAD51-dependent DNA interstrand cross-link repair. Science 333, 84–87.

Lu, R., O’Rourke, J.J., Sobinoff, A.P., Allen, J.A.M., Nelson, C.B., Tomlinson, C.G., Lee, M., Reddel, R.R., Deans, A.J., and Pickett, H.A. (2019). The FANCM-BLM-TOP3A-RMI complex suppresses alternative lengthening of telomeres (ALT). Nat Commun l0, 2252.

Meetei, A.R., de Winter, J.P., Medhurst, A.L., Wallisch, M., Waisfisz, Q., van de Vrugt, H.J., Oostra, A.B., Yan, Z., Ling, C., Bishop, C.E., et al. (2003). A novel ubiquitin ligase is deficient in Fanconi anemia. Nature genetics 35, 165–170.

Meetei, A.R., Medhurst, A.L., Ling, C., Xue, Y., Singh, T.R., Bier, P., Steltenpool, J., Stone, S., Dokal, I., Mathew, C.G., et al. (2005). A human ortholog of archaeal DNA repair protein Hef is defective in Fanconi anemia complementation group M. Nature genetics 37, 958–963.

Menghi, F., Barthel, F.P., Yadav, V., Tang, M., Ji, B., Tang, Z., Carter, G.W., Ruan, Y., Scully, R., Verhaak, R.G.W., et al. (2018). The Tandem Duplicator Phenotype Is a Prevalent Genome-Wide Cancer Configuration Driven by Distinct Gene Mutations. Cancer Cell 34, 197–210 e195.

Menghi, F., Inaki, K., Woo, X., Kumar, P.A., Grzeda, K.R., Malhotra, A., Yadav, V., Kim, H., Marquez, E.J., Ucar, D., et al. (2016). The tandem duplicator phenotype as a distinct genomic configuration in cancer. Proc Natl Acad Sci U S A 113, E2373–2382.

Miki, Y., Swensen, J., Shattuck-Eidens, D., Futreal, P.A., Harshman, K., Tavtigian, S., Liu, Q., Cochran, C., Bennett, L.M., Ding, W., et al. (1994). A strong candidate for the breast and ovarian cancer susceptibility gene BRCA1. Science 266, 66–71.

Murai, J., Huang, S.Y., Das, B.B., Renaud, A., Zhang, Y., Doroshow, J.H., Ji, J., Takeda, S., and Pommier, Y. (2012). Trapping of PARP1 and PARP2 by Clinical PARP Inhibitors. Cancer research 72, 5588–5599.

Nandi, S., and Whitby, M.C. (2012). The ATPase activity of Fml1 is essential for its roles in homologous recombination and DNA repair. Nucleic Acids Res 40, 9584–9595.

Neelsen, K.J., and Lopes, M. (2015). Replication fork reversal in eukaryotes: from dead end to dynamic response. Nat Rev Mol Cell Biol 16, 207–220.

Neidhardt, G., Hauke, J., Ramser, J., Groß, E., Gehrig, A., Müller, C.R., Kahlert, A.K., Hackmann, K., Honisch, E., Niederacher, D., et al. (2017). Association Between Loss-of-Function Mutations Within the FANCM Gene and Early-Onset Familial Breast Cancer. JAMA oncology 3, 1245–1248.

Nguyen, M.O., Jalan, M., Morrow, C.A., Osman, F., and Whitby, M.C. (2015). Recombination occurs within minutes of replication blockage by RTS1 producing restarted forks that are prone to collapse. eLife 4, e04539.

Nik-Zainal, S., Davies, H., Staaf, J., Ramakrishna, M., Glodzik, D., Zou, X., Martincorena, I., Alexandrov, L.B., Martin, S., Wedge, D.C., et al. (2016). Landscape of somatic mutations in 560 breast cancer whole-genome sequences. Nature 534, 47–54.

Niraj, J., Färkkilä, A., and D’Andrea, A.D. (2019). The Fanconi Anemia Pathway in Cancer. Annual review of cancer biology 3, 457–478.

Pace, P., Mosedale, G., Hodskinson, M.R., Rosado, I.V., Sivasubramaniam, M., and Patel, K.J. (2010). Ku70 corrupts DNA repair in the absence of the Fanconi anemia pathway. Science 329, 219–223.

Pan, X., Drosopoulos, W.C., Sethi, L., Madireddy, A., Schildkraut, C.L., and Zhang, D. (2017). FANCM, BRCA1, and BLM cooperatively resolve the replication stress at the ALT telomeres. Proc Natl Acad Sci U S A 114, E5940–e5949.

Paques, F., and Haber, J.E. (1999). Multiple pathways of recombination induced by doublestrand breaks in Saccharomyces cerevisiae. Microbiology and molecular biology reviews: MMBR 63, 349–404.

Peterlongo, P., Catucci, I., Colombo, M., Caleca, L., Mucaki, E., Bogliolo, M., Marin, M., Damiola, F., Bernard, L., Pensotti, V., et al. (2015). FANCM c.5791C>T nonsense mutation (rs144567652) induces exon skipping, affects DNA repair activity and is a familial breast cancer risk factor. Human molecular genetics 24, 5345–5355.

Prakash, R., Krejci, L., Van Komen, S., Anke Schurer, K., Kramer, W., and Sung, P. (2005). Saccharomyces cerevisiae MPH1 gene, required for homologous recombination-mediated mutation avoidance, encodes a 3’ to 5’ DNA helicase. J Biol Chem 280, 7854–7860.

Prakash, R., Satory, D., Dray, E., Papusha, A., Scheller, J., Kramer, W., Krejci, L., Klein, H., Haber, J.E., Sung, P., et al. (2009). Yeast Mph1 helicase dissociates Rad51-made D-loops: implications for crossover control in mitotic recombination. Genes Dev 23, 67–79.

Prakash, R., Zhang, Y., Feng, W., and Jasin, M. (2015). Homologous recombination and human health: the roles of BRCA1, BRCA2, and associated proteins. Cold Spring Harb Perspect Biol 7, a016600.

Quinet, A., Lemaçon, D., and Vindigni, A. (2017). Replication Fork Reversal: Players and Guardians. Mol Cell 68, 830–833.

Raschle, M., Knipscheer, P., Enoiu, M., Angelov, T., Sun, J., Griffith, J.D., Ellenberger, T.E., Scharer, O.D., and Walter, J.C. (2008). Mechanism of replication-coupled DNA interstrand crosslink repair. Cell 134, 969–980.

Rickman, K., and Smogorzewska, A. (2019). Advances in understanding DNA processing and protection at stalled replication forks. The Journal of cell biology 218, 1096–1107.

Romero, N.E., Matson, S.W., and Sekelsky, J. (2016). Biochemical Activities and Genetic Functions of the Drosophila melanogaster Fancm Helicase in DNA Repair. Genetics 204, 531–541.

Rosado, I.V., Langevin, F., Crossan, G.P., Takata, M., and Patel, K.J. (2011). Formaldehyde catabolism is essential in cells deficient for the Fanconi anemia DNA-repair pathway. Nature structural & molecular biology 18, 1432–1434.

Rosado, I.V., Niedzwiedz, W., Alpi, A.F., and Patel, K.J. (2009). The Walker B motif in avian FANCM is required to limit sister chromatid exchanges but is dispensable for DNA crosslink repair. Nucleic Acids Res 37, 4360–4370.

Saini, N., Ramakrishnan, S., Elango, R., Ayyar, S., Zhang, Y., Deem, A., Ira, G., Haber, J.E., Lobachev, K.S., and Malkova, A. (2013). Migrating bubble during break-induced replication drives conservative DNA synthesis. Nature 502, 389–392.

Schlacher, K., Christ, N., Siaud, N., Egashira, A., Wu, H., and Jasin, M. (2011). Double-strand break repair-independent role for BRCA2 in blocking stalled replication fork degradation by MRE11. Cell 145, 529–542.

Scully, R., Panday, A., Elango, R., and Willis, N.A. (2019). DNA double-strand break repairpathway choice in somatic mammalian cells. Nat Rev Mol Cell Biol.

Silva, B., Pentz, R., Figueira, A.M., Arora, R., Lee, Y.W., Hodson, C., Wischnewski, H., Deans, A.J., and Azzalin, C.M. (2019). FANCM limits ALT activity by restricting telomeric replication stress induced by deregulated BLM and R-loops. Nat Commun 10, 2253.

Singh, T.R., Saro, D., Ali, A.M., Zheng, X.F., Du, C.H., Killen, M.W., Sachpatzidis, A., Wahengbam, K., Pierce, A.J., Xiong, Y., et al. (2010). MHF1-MHF2, a histone-fold-containing protein complex, participates in the Fanconi anemia pathway via FANCM. Mol Cell 37, 879–886.

Stafa, A., Donnianni, R.A., Timashev, L.A., Lam, A.F., and Symington, L.S. (2014). Template switching during break-induced replication is promoted by the Mph1 helicase in Saccharomyces cerevisiae. Genetics 196, 1017–1028.

Sun, W., Nandi, S., Osman, F., Ahn, J.S., Jakovleska, J., Lorenz, A., and Whitby, M.C. (2008). The FANCM ortholog Fml1 promotes recombination at stalled replication forks and limits crossing over during DNA double-strand break repair. Mol Cell 32, 118–128.

Tao, Y., Jin, C., Li, X., Qi, S., Chu, L., Niu, L., Yao, X., and Teng, M. (2012). The structure of the FANCM-MHF complex reveals physical features for functional assembly. Nat Commun 3, 782.

Taylor, A.M.R., Rothblum-Oviatt, C., Ellis, N.A., Hickson, I.D., Meyer, S., Crawford, T.O., Smogorzewska, A., Pietrucha, B., Weemaes, C., and Stewart, G.S. (2019). Chromosome instability syndromes. Nature reviews Disease primers 5, 64.

Whitby, M.C. (2010). The FANCM family of DNA helicases/translocases. DNA Repair (Amst) 9, 224–236.

Willis, N.A., Chandramouly, G., Huang, B., Kwok, A., Follonier, C., Deng, C., and Scully, R. (2014). BRCA1 controls homologous recombination at Tus/Ter-stalled mammalian replication forks. Nature 510, 556–559.

Willis, N.A., Frock, R.L., Menghi, F., Duffey, E.E., Panday, A., Camacho, V., Hasty, E.P., Liu, E.T., Alt, F.W., and Scully, R. (2017). Mechanism of tandem duplication formation in BRCA1-mutant cells. Nature 551, 590–595.

Willis, N.A., Panday, A., Duffey, E.E., and Scully, R. (2018). Rad51 recruitment and exclusion of non-homologous end joining during homologous recombination at a Tus/Ter mammalian replication fork barrier. PLoS genetics 14, e1007486.

Willis, N.A., Rass, E., and Scully, R. (2015). Deciphering the Code of the Cancer Genome: Mechanisms of Chromosome Rearrangement. Trends Cancer 1, 217–230.

Willis, N.A., and Scully, R. (2016). Spatial separation of replisome arrest sites influences homologous recombination quality at a Tus/Ter-mediated replication fork barrier. Cell Cycle 15, 1812–1820.

Wu, L., and Hickson, I.D. (2003). The Bloom’s syndrome helicase suppresses crossing over during homologous recombination. Nature 426, 870–874.

Xu, X., Wagner, K.U., Larson, D., Weaver, Z., Li, C., Ried, T., Hennighausen, L., Wynshaw-Boris, A., and Deng, C.X. (1999a). Conditional mutation of Brca1 in mammary epithelial cells results in blunted ductal morphogenesis and tumour formation. Nature genetics 22, 37–43.

Xu, X., Weaver, Z., Linke, S.P., Li, C., Gotay, J., Wang, X.W., Harris, C.C., Ried, T., and Deng, C.X. (1999b). Centrosome amplification and a defective G2-M cell cycle checkpoint induce genetic instability in BRCA1 exon 11 isoform-deficient cells. Mol Cell 3, 389–395.

Xue, X., Sung, P., and Zhao, X. (2015). Functions and regulation of the multitasking FANCM family of DNA motor proteins. Genes Dev 29, 1777–1788.

Xue, Y., Li, Y., Guo, R., Ling, C., and Wang, W. (2008). FANCM of the Fanconi anemia core complex is required for both monoubiquitination and DNA repair. Human molecular genetics 17, 1641–1652.

Yan, Z., Delannoy, M., Ling, C., Daee, D., Osman, F., Muniandy, P.A., Shen, X., Oostra, A.B., Du, H., Steltenpool, J., et al. (2010). A histone-fold complex and FANCM form a conserved DNA-remodeling complex to maintain genome stability. Mol Cell 37, 865–878.

Zeman, M.K., and Cimprich, K.A. (2014). Causes and consequences of replication stress. Nat Cell Biol 16, 2–9.

Zhang, J., and Walter, J.C. (2014). Mechanism and regulation of incisions during DNA interstrand cross-link repair. DNA Repair (Amst) 19, 135–142.

Zheng, X.F., Prakash, R., Saro, D., Longerich, S., Niu, H., and Sung, P. (2011). Processing of DNA structures via DNA unwinding and branch migration by the S. cerevisiae Mph1 protein. DNA Repair (Amst) 10, 1034–1043.

